# A previously-unrecognized motif of transcription factor RYBP, hotspot of cancer-related mutations, is essential for the integrity of *Polycomb* repressive complex 1

**DOI:** 10.1101/2023.10.23.563594

**Authors:** Catarina S. Silva, Laura Mariño Pérez, Irene Garcia Ferrer, Ines Dieryck, Ombeline Pessey, Elisabetta Boeri Erba, Malene Ringkjøbing Jensen, Marco Marcia

## Abstract

*Polycomb* repressive complex 1 (PRC1) catalyzes monoubiquitination of histone H2A on Lys119, promoting gene silencing. Cells at different developmental stages and in different tissues express different PRC1 isoforms. All isoforms share the same catalytic core (subunits RING1B and PCGF) and vary in the composition of regulatory subunits, clustering in two major classes. Canonical isoforms (cPRC1) are regulated by CBX-like subunits, while variant isoforms (vPRC1) are regulated by RYBP-like subunits. The molecular bases for how regulatory subunits affect the structural assembly of the complex and its catalytic activity are still largely unknown. To fill this knowledge gap, here we have specifically studied how RYBP regulates vPRC1 structure and function. Integrating the machine-learning algorithm AlphaFold2 and NMR, we have identified novel vPRC1 structural motifs in RING1B and RYBP. While the new RING1B motif is dispensable for vPRC1 assembly, the RYBP motif is essential for mediating inter-subunit interactions between RYBP and the catalytic RING1B-PCGF4 heterodimer. Importantly, the RYBP motif harbors cancer-related mutations systematically positioned on the same face of a putative transiently-forming α-helix. Biochemical, biophysical and enzymatic characterization of purified cancer-related mutants confirm that this region is crucial for the structural stability of the complex. Overall, our data offer novel insights into the molecular architecture of vPRC1 and the effects of its regulatory subunit on the biochemical, structural, enzymatic, and physio-pathological properties of the complex.

## INTRODUCTION

The evolutionarily conserved *Polycomb* repressive complexes 1 (PRC1) and 2 (PRC2) function as histone remodeling enzymes to synergistically silence gene expression during embryonic development, stem cell differentiation, cell fate determination and cell cycle control (Aloia et al., 2013; Chan and Morey, 2019; Schuettengruber et al., 2017; Simon and Kingston, 2009). Their dysregulation thus frequently associates to cancer or developmental disorders (Chan and Morey, 2019; Flavahan et al., 2017; Pasini and Di Croce, 2016; Schuettengruber et al., 2017).

While PRC2 is recruited to chromatin also *via* PRC1-independent pathways (Healy and Bracken, 2020), the most prominent mechanistic model that rationalizes *Polycomb*-dependent gene silencing suggests that PRC2 recognizes H2AK119Ub1 marks deposited by certain PRC1 isoforms (named variant PRC1, or vPRC1) and deposits H3K27me3 marks, which are in turn recognized by other PRC1 isoforms (named canonical PRC1, or cPRC1) (Blackledge et al., 2014; Cooper et al., 2014; Wang et al., 2004). This model confers vPRC1 isoforms a key role in controlling gene expression and indeed knock-outs of various vPRC1-specific subunits in mice cause early embryonic lethality (Bajusz et al., 2018; Blackledge et al., 2014; Endoh et al., 2017), whereas knock-outs of cPRC1-specific subunits have milder or delayed defects on mice development (Bajusz et al., 2018; Lau et al., 2017).

vPRC1 isoforms are specifically defined as PRC1 complexes whose minimal functional core comprises E3 ubiquitin ligase subunit RING1B (or its paralog RING1A), one of six *Polycomb* group ring-finger domain (PCGF) subunits, and the transcription factor RING1 and YY1-binding protein (RYBP), or its paralog YY1 associated factor 2 (YAF2) (Gao et al., 2012; Tavares et al., 2012). RYBP, which is specifically present in vPRC1 but not in cPRC1 isoforms, is a critical subunit because it enhances the enzymatic activity of the catalytic RING1B/PCGF heterodimer (Blackledge et al., 2014; Fursova et al., 2019; Gao et al., 2012; Rose et al., 2016; Taherbhoy et al., 2015). Besides regulating *Polycomb*-dependent gene silencing, RYBP also participates in various PRC1-independent cellular pathways, for example through its role in apoptosis. Indeed, RYBP has been found to interact with several death effector domain (DED)-containing proteins, as is the case of FADD, procaspase 8 and 10 (in the cytoplasm) and DEDD (in the nucleus), enhancing CD95- and DEDD-mediated apoptosis, respectively. Furthermore, RYBP has been shown to directly interact with apoptin in the nucleus of tumor cells, inducing their apoptosis; to interact and up-regulate FANK1 in tumor cells, inducing apoptosis via the JNK-AP1 signaling pathway; and to bind to and inhibit the function of the E3 ubiquitin ligase MDM2 in the proteosomal degradation of p53, linking RYBP with tumor-associated apoptosis (Simoes da Silva et al., 2018).

Despite its critical role in regulating gene expression, however, RYBP is still very poorly characterized at the molecular level, partly due to its intrinsic structural disorder (Neira et al., 2009). RYBP comprises an N-terminal Npl4 (RanBP2) Zn-finger domain, responsible for interacting with ubiquitin and putatively DNA, a central Lys-rich region of yet unknown function, a C-terminal β-hairpin motif, which interacts with RING1B, and a C-terminal Ser-/Thr-enriched region, also still functionally uncharacterized (Arrigoni et al., 2006; Bejarano et al., 2005; Neira et al., 2009). Only the β-hairpin and N-terminal Zn-finger (through homology to the YAF2 Zn-finger) motifs are structurally resolved (Wang et al., 2010), providing a still largely incomplete picture of the RYBP molecular architecture, necessary to understand its gene regulatory functions at the molecular level.

To provide new insights into its regulatory catalytic function, here we have set out to study how RYBP interacts with the catalytic core of vPRC1 isoforms. Using the machine-learning algorithm AlphaFold2, we have identified a previously-unrecognized structural motif, between the RYBP Zn-finger domain and its Lys-rich core region, where a peculiar pattern of cancer-related mutations occurs. Using NMR spectroscopy coupled to limited proteolysis, crosslinking mass spectrometry (XL-MS), evolutionary alignments and enzymatic data, we have established that the AlphaFold2-identified motif is essential for vPRC1 assembly. Our data thus provide a biochemical rationale to understand the mechanism of severe cancer-relevant mutations in RYBP.

## RESULTS

### AlphaFold2 predicts two previously-unrecognized structural motifs in RYBP and RING1B

Considering that full-length experimental structures of vPRC1 are still missing, we have used the machine-learning algorithm AlphaFold2 to obtain a model of each one of its core subunits in isolation (RING1B, PCGF4, and RYBP), and to attempt predicting the structure of the trimer (Jumper et al., 2021). For this purpose, we submitted relevant sequences (**Table S1**) to the AlphaFold2 colab notebook (Mirdita et al., 2022) and compared the five resulting models obtained for each subunit, based on their pIDDT (predicted local distance difference test) and PAE (predicted alignment error). While the prediction overall confirmed the previously-estimated domain organization of the three vPRC1 subunits, AlphaFold2 predicted two new structural motifs, one in RING1B and one in RYBP (**Figure 1 and Supp. Figure 1**).

**Figure 1:**
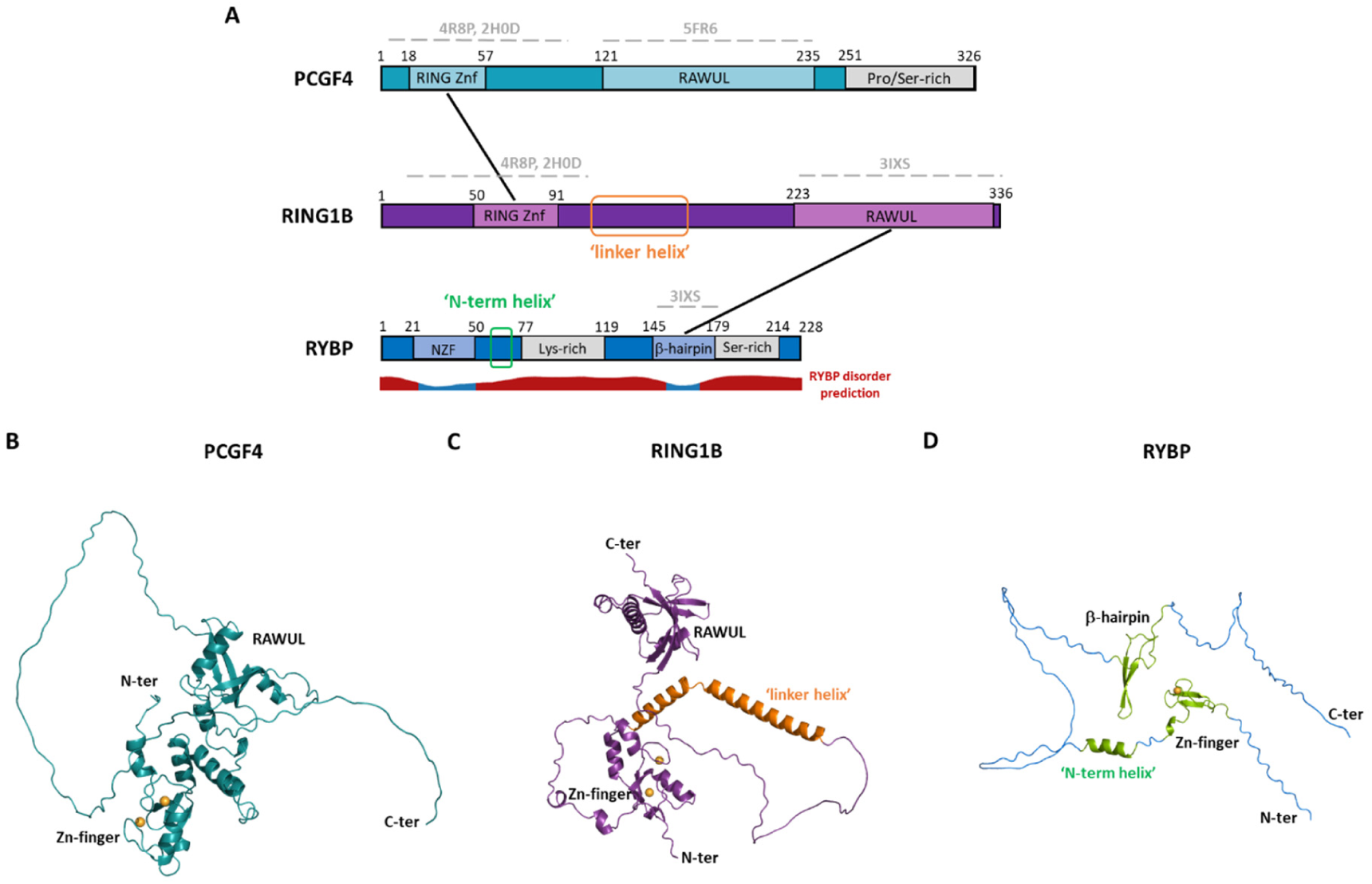
vPRC1.4 core complex subunits and positioning of the two new structural motifs identified by AlphaFold2. **(A)** Schematic representation of the subunits forming the vPRC1.4 core complex. Regions of known structure are indicated by dashed grey lines above the corresponding subunits portions and the corresponding PDB IDs indicated. Known domains and regions are highlighted and reported interactions between domains indicated by black lines. The two new structural motifs predicted by AlphaFold2 are indicated in orange and green. RYBP disorder prediction is indicated below RYBP subunit. The figure is drawn to scale based on the length of each subunit. AlphaFold2 prediction models for **(B)** PCGF4, **(C)** RING1B and **(D)** RYBP subunits in cartoon representation. Structured known domains are indicated as well as the two new structural motifs identified by AlphaFold2, RING1B ‘linker helix’ in orange and RYBP ‘N-term helix’ in green.

In RING1B, the newly-identified structured region is a putative helix-loop-helix motif (hereafter called RING1B ‘linker helix’) that localizes in the core region of the subunit, between its RING and RAWUL domains, and that was previously predicted to be unstructured. This RING1B ‘linker helix’ spans amino acids 117-165 with the loop covering residues 135-136. The likelihood of the prediction of such motif is corroborated by a high pIDDT score for the majority of the helix (above 0.7 until residue R158, with an average value of 0.86, then below 0.7 with an average value of 0.55 for residues 159-165).

In RYBP, the newly identified structural motif is a putative α-helix that comprises amino acids 59-69 (hereafter called RYBP ‘N-term helix’), which are positioned just after the N-terminal Zn-finger and were expected to be unstructured to date. The likelihood of the prediction is corroborated by the pIDDT score (mostly above 0.7, with some variability between models, with an average value of 0.72).

Besides these novel motifs, the known RING Zn-finger and RAWUL domains of RING1B and PCGF4, and the RYBP Zn-finger and β-hairpin are the only other structured regions predicted for vPRC1.

Finally, the AlphaFold2 prediction for putative inter-subunit interactions, in the context of trimeric vPRC1, confirmed the known interaction between the RING1B and PCGF4 Zn-finger domains. Moreover, when attempting to predict the structure of the trimer, both the RING1B ‘linker helix’ and the RYBP ‘N-term helix’ were systematically predicted in the same conformations as for the individual subunits. But AlphaFold2 failed to identify any other confident inter-subunit interaction in vPRC1, suggesting high flexibility of this complex (**Supp. Figure 1**).

Overall, although it fails to predict the complete assembly of the trimeric vPRC1 complex, AlphaFold2 predicts two potentially novel structural motifs, the RING1B ‘linker helix’ and the RYBP ‘N-term helix’.

### The newly-predicted motifs are evolutionarily conserved

To gain further evidence on the formation and relevance of the newly-predicted RING1B ‘linker helix’ and the RYBP ‘N-term helix’, we then performed evolutionary alignments to assess the degree of conservation of these specific motifs (**Figure 2**).

**Figure 2:**
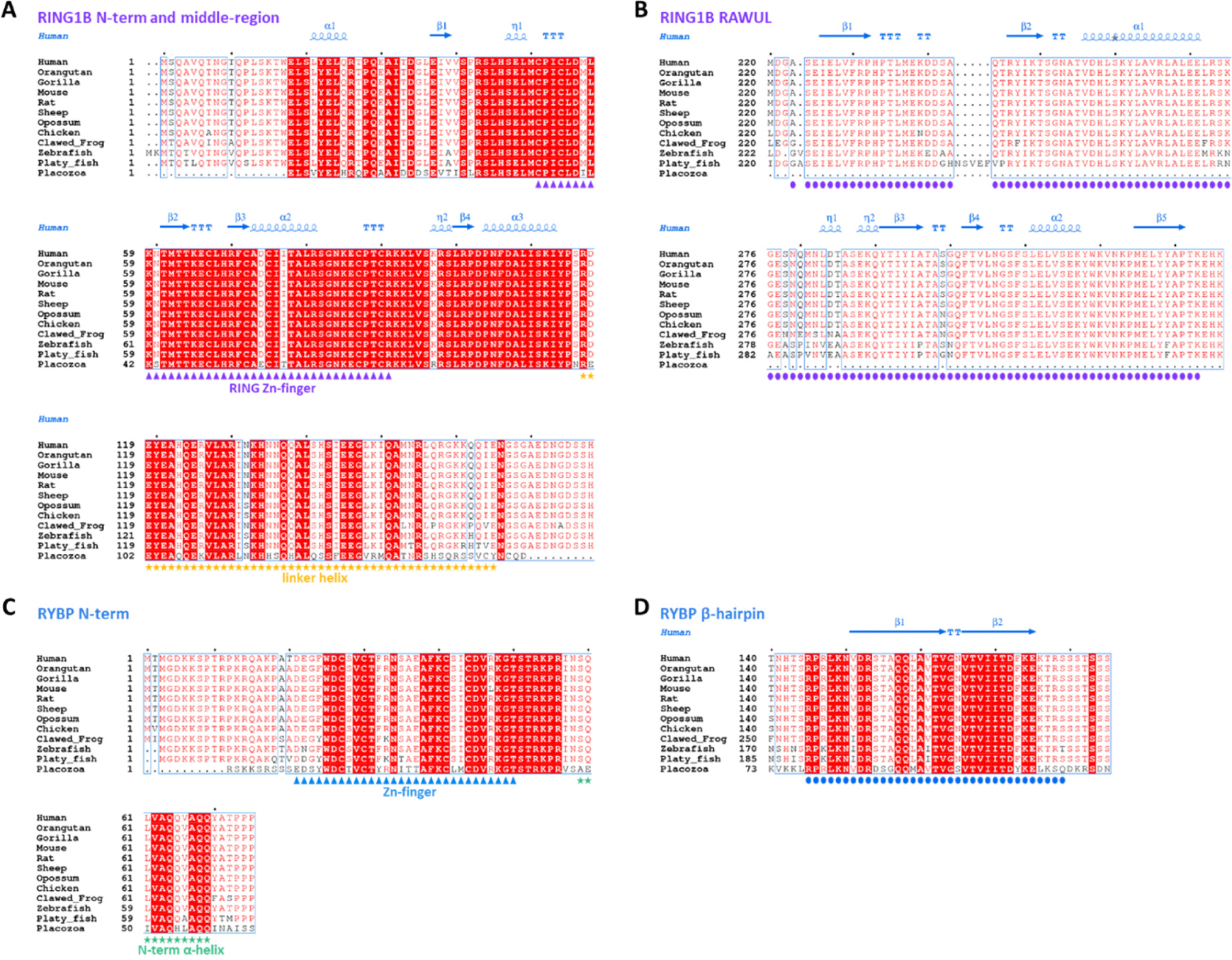
Evolutionary sequence conservation of RING1B ‘linker helix’ and RYBP ‘N-term helix’. Sequence alignment of human (*H. sapiens*), orangutan (*P. abelii*), gorilla (*G. gorilla)*, mouse (*M. musculus*), rat (*R. norvegicus*), sheep (*O. aries*), opossum (*M. domestica*), chicken (*G. gallus*), clawed frog (*X. laevis*), zebrafish (*D. rerio*), platy fish (*X. maculatus*) and placozoa (*T. adhaerens*) RING1B and RYBP subunits. Sequence numbering is shown on the left and black dots above the sequences mark every tenth residue for the first sequence. Highly conserved regions are boxed, with similar residues represented in red against a white background, invariant residues represented against a red background and non-conserved residues indicated in black. **(A)** Aligned RING1B N-terminal and middle linker region amino acid sequences. The secondary structure annotation of human RING1B RING Zn-finger, as derived from its three-dimensional X-ray structure (PDB 2H0D), is depicted in blue on top of the aligned sequences; α-helices are represented by spirals and β-strands by arrows (alpha helices (α); beta-strands (β); 3_10_-helix (η); strict α-turns (TTT); strict β-turns (TT)). Purple triangles indicate RING Zn-finger positioning and yellow stars indicate the newly predicted RING1B ‘linker helix’ structural motif. **(B)** Aligned RING1B RAWUL domain amino acid sequences. The secondary structure annotation of human RING1B RAWUL, as derived from its three-dimensional X-ray structure (PDB 3IXS, chain B), is depicted in blue on top of the aligned sequences (annotations are as per A)). Purple circles indicate RING1B RAWUL domain positioning. **(C)** Aligned RYBP N-terminal region amino acid sequences. Blue triangles indicate RYBP Zn-finger positioning and green stars indicate the newly predicted RYBP ‘N-term helix’ structural motif. **(D)** Aligned RYBP β-hairpin amino acid sequences. The secondary structure annotation of human RYBP β-hairpin, as derived from its three-dimensional X-ray structure (PDB 3IXS, chain A), is depicted in blue on top of the aligned sequences (annotations are as per A)). Blue circles indicate RYBP β-hairpin positioning.

The RING1B ‘linker helix’ is conserved from Primates to basal Metazoa, showing a high degree of sequence conservation until basal Vertebrata. Levels of sequence conservation span from 88% in platy fish to 100% in Primates, with Placozoa (*Trichoplax adhaerens*), the simplest in structure of all animals, displaying approximately 51% of sequence identity and 86% of sequence similarity. Comparison to the level of sequence conservation of RING1B remaining structured regions shows that RING Zn-finger domain is equally highly conserved until basal Metazoa, with nearly 100% of sequence conservation across metazoans, with Placozoa showing ∼83% sequence identity (**Figure 2A**). As for the RING1B RAWUL domain, no equivalent was found in *Trichoplax adhaerens*, despite high level of sequence conservation of this domain among the other species analyzed (mostly 92 to 100% of sequence identity, with zebrafish and platy fish showing levels of 85% and 82%, respectively) (**Figure 2B**).

Similar to the RING1B ‘linker helix’, also the RYBP ‘N-term helix’ is highly conserved among metazoans, with 91 to 100% sequence conservation, except for Placozoa which shows ∼55% of sequence identity, but ∼86% of sequence similarity. The same level of sequence conservation is observed for RYBP N-terminal Zn-finger and C-terminal β-hairpin domains, for which levels of sequence identity span from ∼87 to 100% and from ∼89 to 100%, respectively, except for the Placozoa. In *Trichoplax adhaerens*, despite RYBP having about half the size of the human RYBP, both functional domains are preserved with its Zn-finger displaying ∼63% sequence conservation (and ∼92% sequence similarity) and its β-hairpin ∼71% sequence identity (and ∼93% similarity) (**Figure 2C and 2D**).

Overall, our sequence alignments suggest that the newly-identified structured motifs of RING1B and RYBP, predicted by AlphaFold2, are highly conserved throughout evolution, suggesting that they may represent functionally-important regions of these proteins, possibly connected to their vPRC1-related or to PRC1-unrelated functions.

### vPRC1.4 can be purified to homogeneity in a functionally-active form, and adopts an elongated topology

Encouraged by the high degree of sequence conservation of the newly-predicted RING1B ‘linker helix’ and the RYBP ‘N-term helix’, we then set out to further characterize these motifs experimentally.

Towards such goal, we first heterologously expressed trimeric wild type vPRC1.4, composed of subunits RING1B, PCGF4 and RYBP, in Sf21 insect cells using the MultiBac co-expression system, as we had previously done for cPRC1 isoforms (Colombo et al., 2019). We obtained a pure and homogeneous sample, which assembles into the expected heterotrimer with 1:1:1 stoichiometry of its three subunits, as judged by size exclusion chromatography coupled to multi-angle laser light scattering (SEC-MALLS) and by native mass spectrometry (native MS) (**Figure 3A and 3B**). SEC-MALLS estimated a total molecular weight of 98.6 kDa, with a polydispersity of 1.0, while native-MS a total molecular weight of 99.8 kDa, in line with the theoretical molecular weight of the complex estimated from the sequence of its subunits, 99.6 kDa. By analytical ultracentrifugation (AUC), we further confirmed the homogeneity of vPRC1.4, its experimental molecular weight (97 kDa), and its hydrodynamic properties, i.e. its hydrodynamic radius (R_h_) = 5.14 nm, sedimentation coefficient (s) = 2.83 and frictional ratio (f/f_0_) = 1.69 (**Figure 3C**). These hydrodynamic properties are in line with SAXS data (R_g_ = 5.21 nm, D_max_ = 19.9 nm), which further allow the determination of the vPRC1.4 elongated molecular topology (**Figure 3D-H**).

**Figure 3:**
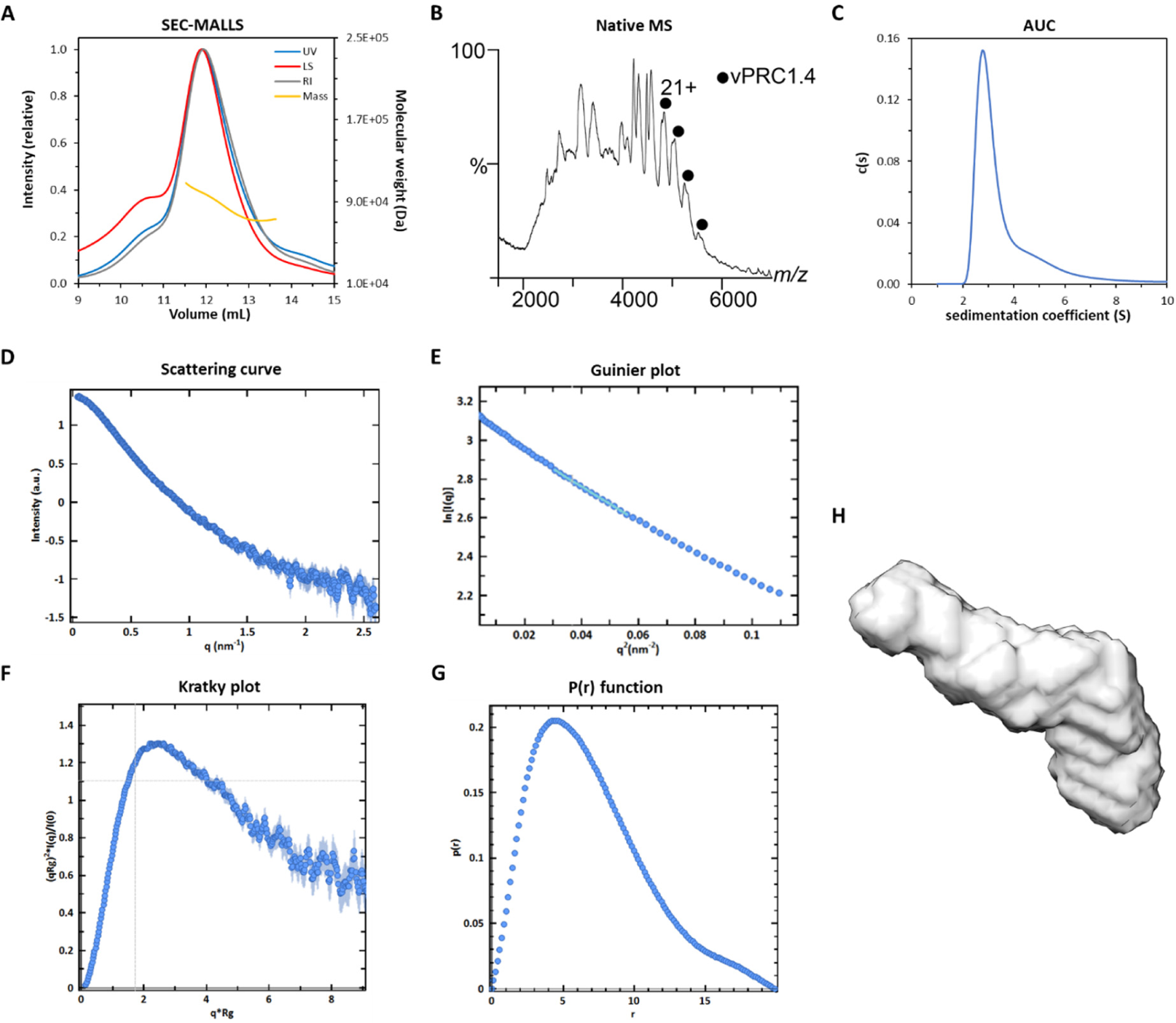
vPRC1.4 core complex purifies to homogeneity and assembles into a stoichiometric and elongated trimeric complex. Biophysical characterization of vPRC1.4 core complex by **(A)** Size exclusion chromatography/multi-angle laser light scattering (SEC-MALLS), **(B)** Native-MS and **(C)** Analytical ultracentrifugation (AUC) indicate a homogeneous sample and a complex stoichiometry of 1:1:1. **(D-H)** SAXS experiments on vPRC1.4. **(D)** Scattering curve for the integrated frames. **(E)** Close-up of the Guinier region. The linear fit is shown in cyan. **(F)** Normalized Kratky plot. The shape of the curve is indicative of a flexible particle with the grey crosshair indicating the position of the peak for an expected globular protein. **(G)** P(r) distribution function **(H)** Bead volume obtained for vPRC1.4 core complex reveals an elongated shape.

Importantly, the reconstituted wild type vPRC1.4 core complex is capable of binding and monoubiquitinating nucleosomes. First, size exclusion chromatography and SDS-PAGE electrophoresis show that isolated vPRC1.4 elutes with a retention volume ∼1.38 mL, reconstituted nucleosome core particles (NCPs) elute with a retention volume ∼1.24 mL, and the complex formed by vPRC1.4 with NCPs elutes with a retention volume ∼1.15 mL from a Superdex 200 increase 5/150 GL column, compatible with complex formation (**Figure 4A and 4B**). Electrophoretic mobility shift assays (EMSA) further confirmed the binding of vPRC1.4 to NCPs (**Figure 4C**). Second, H2A monoubiquitination activity assays showed that our isolated vPRC1.4 core complex is functionally active (**Figure 4D**).

**Figure 4:**
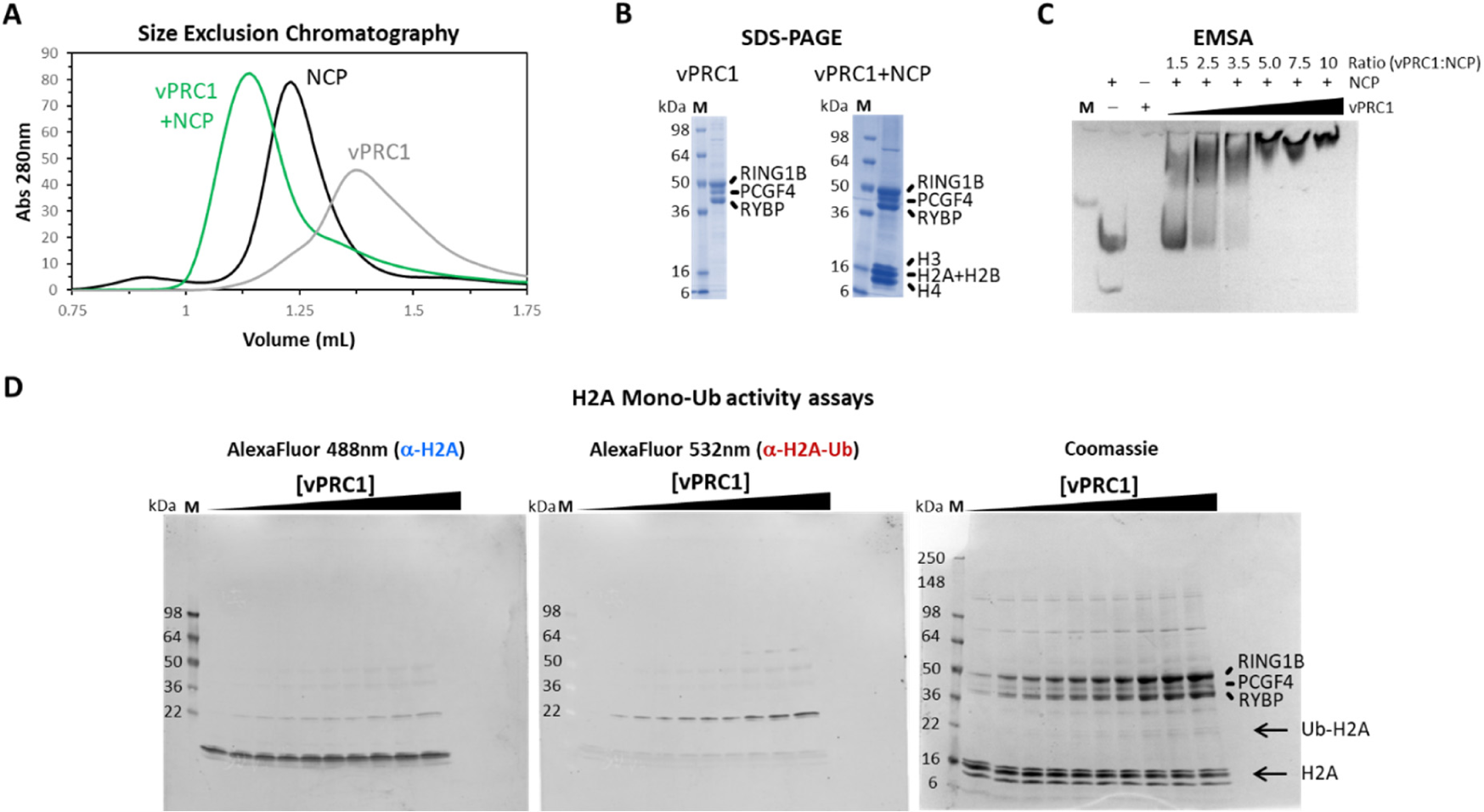
vPRC1.4 core complex is functionally active, and thus capable of binding and monoubiquitinating nucleosomes. **(A)** Size exclusion chromatography (SEC) elution profiles obtained for vPRC1.4 (grey) and nucleosomes (NCPs; black) in isolation, as well as for the vPRC1.4 in complex with NCPs (green) using a S200 increase 5/150GL column. **(B)** SDS-PAGE on final samples of vPRC1.4 in isolation and in complex with NCPs, after SEC. **(C)** Electrophoretic mobility shift assays (EMSA) showing association of vPRC1.4 trimeric complex to NCPs, as vPRC1.4 concentration increases. Gel is stained with Sybr Safe. **(D)** H2A monoubiquitination activity assays on vPRC1.4 core complex show an active complex, as revealed by western blot.

### While the RING1B ‘linker helix’ is dispensable, the RYBP ‘N-term helix’ is resistant to proteolysis within the full-length, *in vitro* purified vPRC1

To better characterize the topology of the isolated vPRC1.4 complex, we employed crosslinking mass spectrometry (XL-MS) using DSS as an alkylating agent (**Figure 5A**). Our data reveal an extensive network of inter-subunit interactions within vPRC1.4. For instance, our data recapitulated the known inter-subunit interaction between RING1B RAWUL and RYBP β-hairpin C-terminal domains, derived from available crystal structures of small vPRC1 subdomains (Wang et al., 2010). Importantly, though, our data also suggest the presence of previously-unrecognized interactions. First, both RING1B Zn-finger and RAWUL domains crosslink with the RYBP N-terminal region, mostly in close proximity to the RYBP ‘N-term helix’. Second, we could also identify novel interactions between the PCGF4 RAWUL domain (in particular its C-terminal region) and both RYBP and RING1B Zn-fingers, between the PCGF4 RAWUL domain and the RYBP β-hairpin, between RYBP Zn-finger and the PCGF4 Zn-finger and Pro/Ser-rich regions. Taken together these crosslinks suggest that, despite its intrinsic disorder, RYBP – and particularly the motifs surrounding the RYBP ‘N-term helix’ – establish extensive interactions with the vPRC1 catalytic core (**Figure 5A**). In addition, a novel inter-subunit interaction was also identified between the RING1B ‘linker helix’ and the C-terminal region of PCGF4 RAWUL domain.

**Figure 5:**
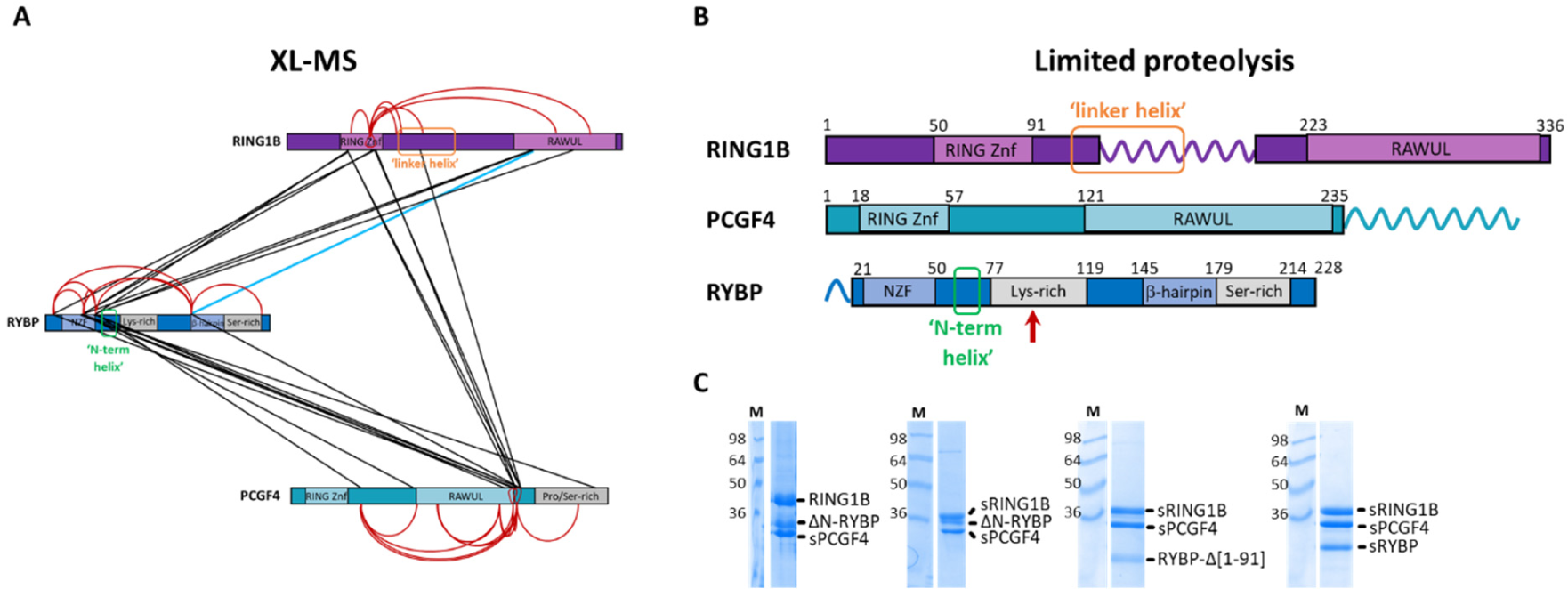
RYBP is resistant to proteolysis within the complex and establishes extensive inter-subunit interactions with vPRC1.4 catalytic dimer, in particular through its N-terminal region. **(A)** Graphical summary illustrating crosslinking mass spectrometry (XL-MS) analysis for the vPRC1.4 complex. Red and black lines indicate intra-subunit and inter-subunit interactions, respectively. The blue line highlights the recapitulated known inter-subunit interaction between RING1B RAWUL and RYBP β-hairpin C-terminal domains. Extensive inter-subunit interactions are established between RYBP and vPRC1.4 catalytic dimer, particularly through the motifs surrounding RYBP ‘N-term helix’. **(B)** Graphical summary illustrating the results from limited proteolysis. Labile proteolyzed regions are indicated by a spiral. The red arrow indicates a point of proteolysis in RYBP. Despite its intrinsic disorder, RYBP is resistant to proteolysis within the trimer. Very few regions of the complex were sensitive to proteolysis. **(C)** vPRC1.4 core complex re-engineering based on XL-MS and limited proteolysis results. While RING1B ‘linker helix’ is not essential for the trimeric complex assembly, RYBP ‘N-term helix’ might play an important role in complex assembly, as its removal disrupts the complex stoichiometry. sPCGF4 indicates removal of PCGF4 C-terminal region (residues 240-326); ΔN-RYBP corresponds to the deletion of RYBP N-terminus (residues 1-15); sRING1B indicates substitution of RING1B middle region (residues 127-196) for a linker; and RYBP-Δ[1-91] denotes removal of the entire RYBP N-terminus until residue 91, whereas sRYBP indicates simultaneous removal of the RYBP N-terminal amino acids (residues 1-15) and Lys-rich region (residues 71-119).

Having mapped the vPRC1.4 topology, we then employed limited proteolysis using trypsin digestion and peptide mass fingerprinting mass spectrometry (PMF-MS), to identify labile and stable regions of the complex. Surprisingly, despite the expected flexibility of vPRC1.4, also corroborated by the AlphaFold2 predictions (see above), very few regions of the complex were sensitive to proteolysis (**Figure 5B**). Notably, one of these proteolysis-sensitive regions covers the majority of RING1B ‘linker helix’, including the region of crosslink between the RING1B ‘linker helix’ and PCGF4 (residues 127-196). Other regions sensitive to proteolysis expectably include the PCGF4 C-terminal Pro/Ser-rich region (residues 228-326), the RYBP N-terminus (residues 1-15, preceding the Zn-finger domain), and amino acid 92 in the RYBP Lys-rich region, downstream of the RYBP ‘N-term helix’ (proteolysis at amino acid residue 92). However, the RYBP ‘N-term helix’ itself is resistant to proteolysis.

On the basis of the XL-MS and limited proteolysis data and to specifically test the contribution of the RING1B ‘linker helix’, PCGF4 C-terminal Pro/Ser-rich region and RYBP ‘N-term helix’ to the overall assembly of vPRC1, we have then engineered the complex by removing selected domains. First, removal of PCGF4 C-terminal region (residues 240-326; sPCGF4) and of RYBP N-terminus (residues 1-15; ΔN-RYBP), yields a stochiometric trimeric complex. Second, further substitution of RING1B middle region (residues 127-196, containing RING1B ‘linker helix’), with a shorter serine-glycine linker, revealed that this intermediary region is not essential for maintaining the complex together (sRING1B). Third, removal of the RYBP N-terminus (Zn-finger and ‘N-term helix’, i.e. residues 1-91; RYBP-Δ[1-91]) yields a non-stoichiometric complex (**Figure 5C**). Instead, simultaneous removal of the RYBP N-terminal amino acids (residues 1-15) and Lys-rich region (residues 71-119) (sRYBP) results in stoichiometric trimeric complex formation (**Figure 5C**).

Taken together these data suggest that (i) the RING1B ‘linker helix’ is unlikely to be involved in assembly or stabilization of vPRC1, because it does not seem essential for the complex to form and readily degrades by proteolysis; (ii) RYBP is overall unexpectedly stable within the trimer, probably thanks to the establishment of many inter-subunit contacts between loosely structured RYBP motifs and the catalytic vPRC1 core which cannot be modelled by AlphaFold2; and (iii) within RYBP, the ‘N-term helix’ is particularly stable, close to a densely-interacting region of the trimer, and possibly important for complex assembly, because removing the RYBP N-terminus where this putative helix is located disrupts the vPRC1 stoichiometry.

### The RYBP ‘N-term helix’ interacts with the RING1B-PCGF4 heterodimer

Established that the newly-identified RYBP ‘N-term helix’ is resistant to proteolysis and close to sites of inter-subunit interaction, we focused on this region more closely, and used nuclear magnetic resonance (NMR) to characterize its contribution to the assembly of RYBP with the catalytic RING1B-PCGF4 heterodimer. We set out to first study RYBP in isolation, then in the context of the full-length trimer, and finally in complex with individual domains of RING1B and PCGF4.

To determine the structural propensities of RYBP in isolation, we expressed this subunit in *E. coli*, and successfully isolated pure, homogeneous, ^13^C/^15^N-labelled RYBP for NMR studies (**Supp. Figure 2A-B**). A series of triple resonance experiments allowed for spectral assignment of the ^1^H-^15^N HSQC spectrum (**Figure 6A**), except for the residues belonging to the Zn-finger for which magnetization transfer was not complete during the triple resonance experiments. The experimental ^13^Cα and ^13^Cβ chemical shifts were used to derive the secondary structure propensities (**Figure 6B**) showing that, in the free form, RYBP is intrinsically disordered devoid of significant amounts of secondary structure with only its N-terminal Zn-finger being partly structured (residues 21-53), as evidenced from the dispersion of the NMR signals in the ^1^H-^15^N HSQC spectrum (**Figure 6A**). Specifically, in the context of free RYBP, the RYBP β-hairpin is unstructured, while the putative ‘N-term helix’ only shows weak α-helical propensity (20%) (**Figure 6B**).

**Figure 6:**
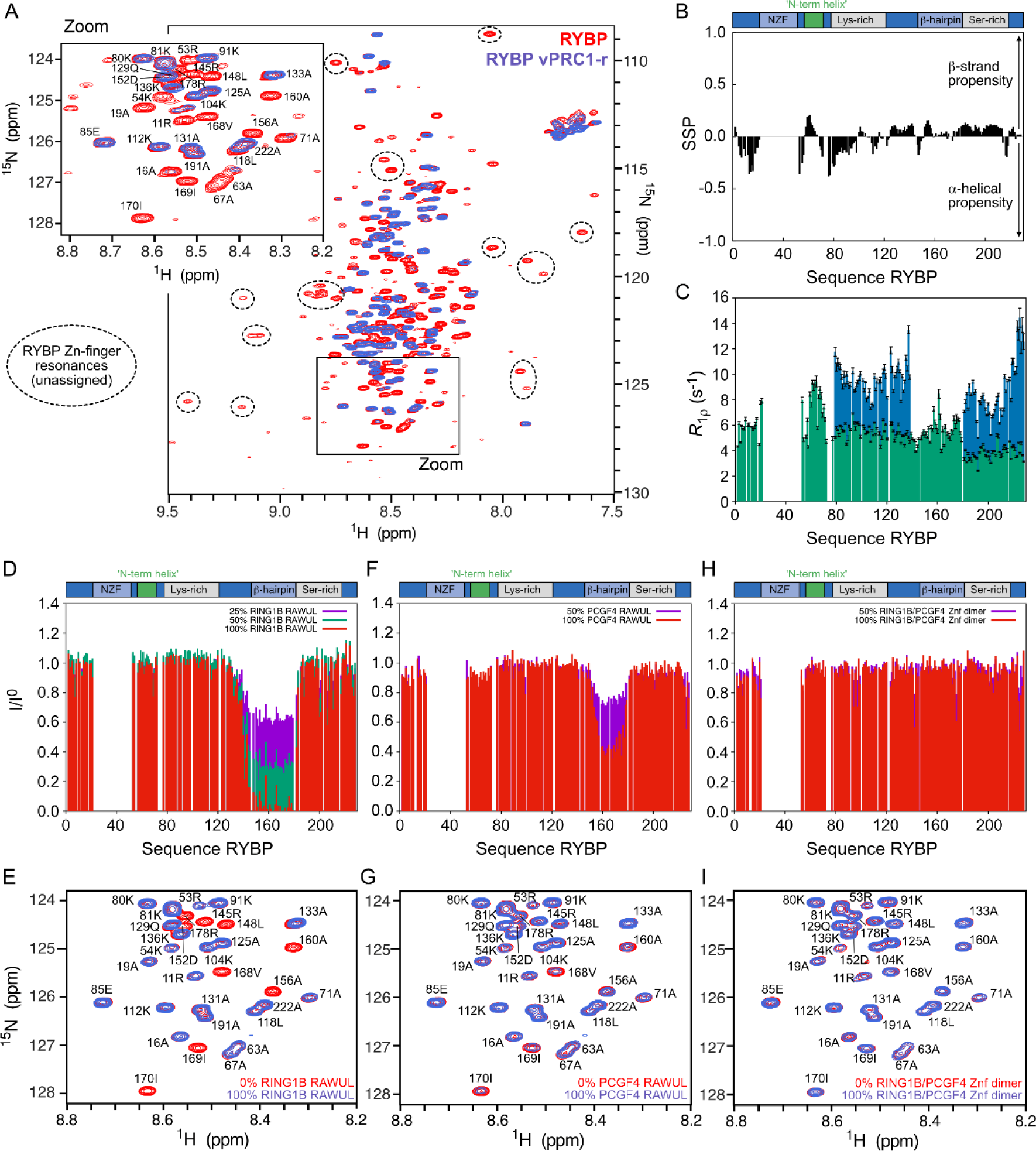
Conformational propensities of RYBP and its interaction with RING1B and PCGF4. **(A)** ^1^H-^15^N HSQC spectrum of RYBP in isolation (red) and superimposed on that of RYBP in the context of the reconstituted trimeric complex (blue). A zoom is shown with labels corresponding to the spectral assignment. Several well-dispersed signals (unassigned and marked with dashed circles) are observed in the ^1^H-^15^N HSQC spectrum that presumably originate from the folded Zn-finger domain. **(B)** Secondary structure propensity (SSP) of free RYBP calculated from the experimental ^13^Cα and ^13^Cβ chemical shifts. The resonances corresponding to the residues of the Zn-finger remain unassigned. **(C)** ^15^N *R*_1ρ_ relaxation rates of free RYBP (green) and of RYBP in the context of the reconstituted trimeric complex (blue). The relaxation rates were measured at 5°C and at a ^1^H frequency of 950 MHz. **(D)** NMR titration experiment of ^15^N-labeled RYBP with increasing concentrations (25, 50 and 100% molar ratio compared to RYBP) of RING1B RAWUL. The peak intensity ratio, I/I^0^, is reported, where I corresponds to the intensities of the resonances in the ^1^H-^15^N HSQC spectra with RING1B RAWUL and I^0^ to the intensities of free RYBP. **(E)** Zoom on a region of the ^1^H-^15^N HSQC spectra of free RYBP (red) and with 100% (molar ratio) of RING1B RAWUL (blue). **(F)** NMR titration experiment of ^15^N-labeled RYBP with increasing concentrations (50 and 100% molar ratio compared to RYBP) of PCGF4 RAWUL. **(G)** Zoom on a region of the ^1^H-^15^N HSQC spectra of free RYBP (red) and with 100% (molar ratio) of PCGF4 RAWUL (blue). **(H)** NMR titration experiment of ^15^N-labeled RYBP with increasing concentrations (50 and 100% molar ratio compared to RYBP) of RING1B/PCGF4 Zn-finger dimer **(I)** Zoom on a region of the ^1^H-^15^N HSQC spectra of free RYBP (red) and with 100% (molar ratio) of RING1B/PCGF4 Zn-finger dimer (blue).

We then co-purified the full-length unlabeled RING1B-PCGF4 catalytic dimer, expressed in insect cells (Colombo et al., 2019), with ^13^C/^15^N-labelled RYBP, expressed in *E. coli* as described above. We could successfully reconstitute a pure, homogeneous, stable and functionally active vPRC1.4 trimer (**Supp. Figure 2A-C**). Analysis of the ^1^H-^15^N HSQC spectrum obtained for RYBP within the reconstituted trimer revealed that the lysine and serine-rich regions of RYBP still remain highly dynamic within the complex with no significant chemical shift perturbations compared to free RYBP (**Figure 6A-C**). Measurements of ^15^N *R*_1ρ_ relaxation in free RYBP and within the reconstituted trimer reveal, however, a slight restriction of motion of the lysine and serine-rich regions in the context of the trimer (**Figure 6C**). Moreover, comparison of RYBP HSQC spectra in isolation *vs* reconstituted trimer revealed that three regions of RYBP participate in complex formation or become rigidified upon interaction (**Figure 6A-C**). These regions correspond to residues 1-21, the RYBP N-terminus preceding the Zn-finger domain, residues 54-77, surrounding the previously-uncharacterized putative ‘N-term helix’, and residues 138-186, surrounding the β-hairpin known to acquire a folded conformation upon interaction with the RING1B C-terminal RAWUL domain (Neira et al., 2009; Wang et al., 2010).

In line with our crosslinking, limited proteolysis, and mutagenesis data, the NMR data shows that the RYBP ‘N-term helix’ becomes restricted in motion in the context of the trimer suggesting that it may represent a novel inter-subunit interaction site between RYBP and RING1B/PCGF4 catalytic dimer.

### Interaction of RYBP with RING1B and PCGF4

To more finely map the interactions of the RYBP ‘N-term helix’ with its partner proteins, we performed NMR titration experiments between ^15^N-labelled RYBP and the Zn-finger and RAWUL domains of RING1B and PCGF4 (expressed in *E. coli* and purified to homogeneity) (**Supp. Figure 2D-F**).

Analysis of the ^1^H-^15^N HSQC spectrum, and the corresponding intensity plot, obtained for the titration of RYBP with the RING1B RAWUL domain, showed that RING1B RAWUL domain interacts with the β-hairpin of RYBP (residues 140-182, **Figure 6D-E** and **Supp. Figure 2G**), recapitulating the already reported interaction between these two domains (Wang et al., 2010). We can also observe an unexpected interaction between the PCGF4 RAWUL domain and the RYBP β-hairpin, albeit of weaker affinity than the interaction with the RING1B RAWUL domain, which we also could capture by XL-MS (residues 151-180, **Figure 6F-G** and **Supp. Figure 2H**). Finally, our NMR titrations showed that the RING1B/PCGF4 RING Zn-finger dimer does not interact with any of the RYBP motifs visible in the NMR spectrum (**Figure 6H-I** and **Supp. Figure 2I**).

These titration studies, which cover all RING1B-PCGF4 structured domains, demonstrate that the RYBP ‘N-term helix’ does not directly interact with any of these regions. The RYBP ‘N-term helix’ though certainly undergoes restriction of motion within the complex being tethered by the other folded domains.

### The ‘N-term helix’ is a hotspot of RYBP cancer-relevant mutations, which destabilize the structural integrity of vPRC1

To connect the structural role of the RYBP ‘N-term helix’ to vPRC1 functions, we scouted available databases to screen if this newly-identified motif is in any way related to human diseases. Indeed, using the cBioPortal, Biomuta and Cosmic databases, we have found that the RYBP ‘N-term helix’ is a hotspot of cancer-related mutations. For instance, mutations S59F, Q60* or Q60H (equivalent to Q58H in YAF2), A63T and Q69* or Q69K (equivalent to Q67K in YAF2) are connected to uterine, central nervous system, stomach, lung, head and neck squamous cell, liver and bladder carcinomas or adenocarcinomas, as well as skin melanomas. Interestingly, as evident from the corresponding amino acid numbering, these mutations occur with a very peculiar pattern along the RYBP ‘N-term helix’, every 3-4 amino acids. Such pattern would confine the mutations all on the same side of the putative RYBP ‘N-term helix’ (**Figure 7A**). We have therefore cloned, expressed, purified, and biochemically and enzymatically characterized three representative mutants (Q60H, A63T, and Q69K) to experimentally assess the effect of these mutations on the vPRC1 structural and/or functional integrity.

**Figure 7:**
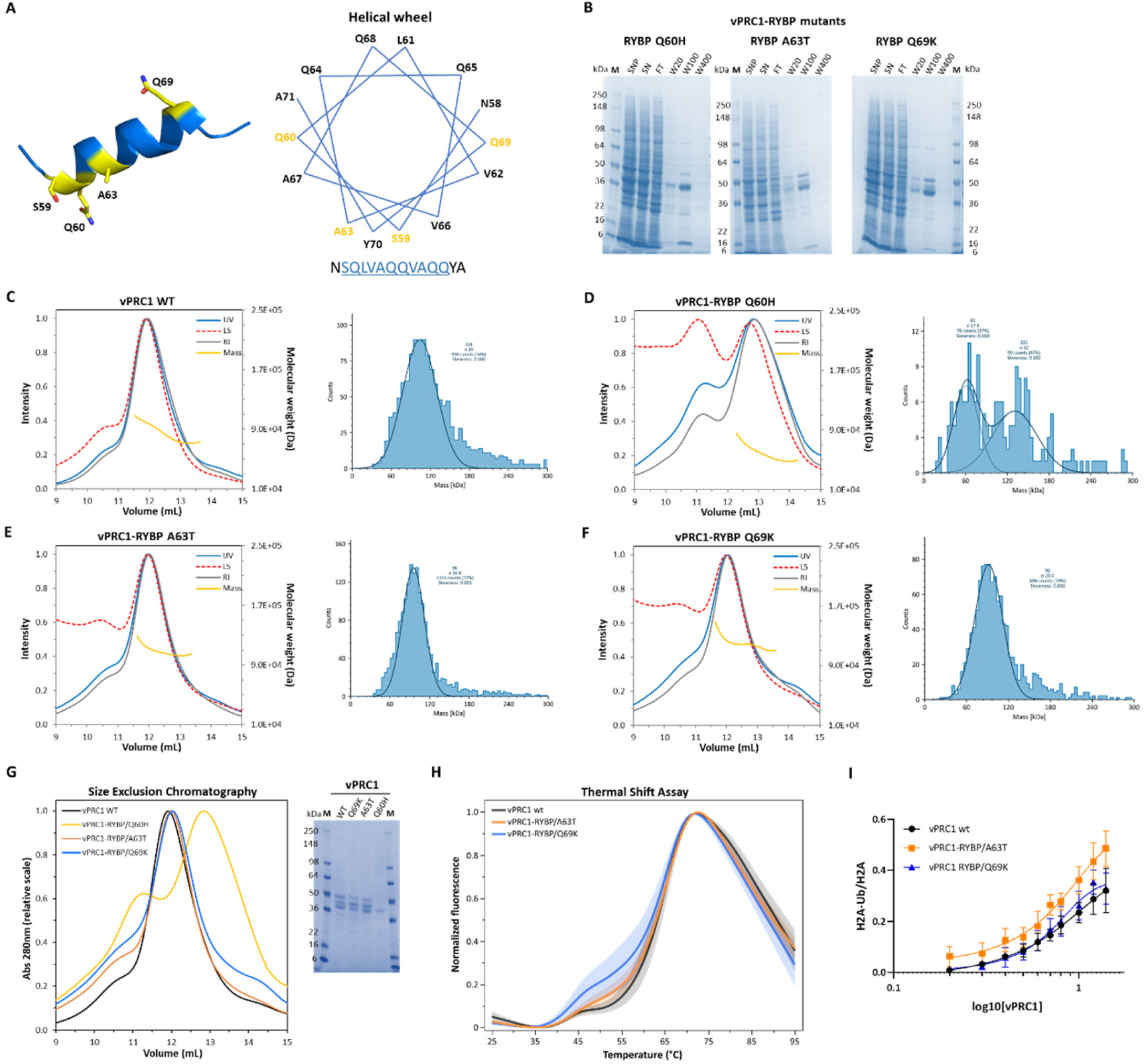
The RYBP ‘N-term helix’ is a hotspot of cancer-relevant mutations which destabilize vPRC1 structural integrity. **(A)** Cartoon representation of RYBP ‘N-term helix’ as predicted by AlphaFold2. Cancer-related mutations found in databases are annotated and seem to follow a peculiar pattern, all sitting on the same side of the putative RYBP ‘N-term helix’, as observed from the helical wheel model. **(B)** Effect of three representative mutants (RYBP Q60H, A63T, and Q69K) on vPRC1 complex integrity, as assessed by Ni-Sepharose pull-downs. Insertion of the RYBP Q60H mutation in vPRC1 disrupts the formation of the trimeric complex. **(C-F)** Biophysical characterization of vPRC1 wt **(C)** and vPRC1-RYBP Q60H **(D)**, A63T **(E)** and Q69K **(F)** mutants by SEC-MALLS and mass photometry (MP). **(G)** Size exclusion chromatography (SEC) elution profiles obtained for vPRC1 wt and vPRC1-RYBP Q60H, A63T and Q69K mutants and respective SDS-PAGE gel on the input samples. **(H)** Thermal denaturation assays performed for vPRC1 wt and vPRC1-RYBP A63T and Q69K mutants. Error bands are constructed using one standard error from the mean. **(I)** Plot reporting the H2A monoubiquitination activities of vPRC1 wt and vPRC1-RYBP A63T and Q69K mutants, as measured by western blot. The errors bars correspond to the standard error of the mean of n=3 independent experiments.

Remarkably, the Q60H mutant induced a dramatic structural destabilization of vPRC1, preventing us from isolating the intact trimer (**Figure 7B and D and Supp. Figure 3**). For the A63T and Q69K mutants, it was instead possible to purify the vPRC1 trimer to homogeneity, as confirmed by SEC-MALLS and mass photometry. These two mutated samples maintained a similar molecular weight as the wild type complex (see above and **Figure 7C, 7E-G and Supp. Figure 3**). However, thermal denaturation assays revealed that with respect to wild type, the two mutants had a different denaturation profile. All three complexes display two melting transitions, at about 43°C (likely corresponding to subunit dissociation and exposure of inter-subunit interacting hydrophobic regions) and at about 63°C (likely corresponding to the complete denaturation of all subunits). However, while for the wild type samples the first transition occurred in only ∼ 11 ± 3 % of the sample, as judged by the intensity of the transition, in the A63T and Q69K mutants inter-subunit dissociation was more pronounced (16 ± 4 % and 25 ± 10 %, respectively, **Figure 7H**), attesting for the higher destabilization effect of the mutants. With respect to wild type, the mutants also showed a statistically significantly different variance across different purifications of the samples (Levene test, p=0.033). Technical replicates of each sample instead were extremely reproducible, with average standard error of the mean values of 1.0%, 0.9% and 1.6% for vPRC1 wt, A63T and Q69K mutants, respectively.

We have further analyzed the effect of the A63T and Q69K mutations on the activity of the isolated vPRC1 trimer *in vitro*. Monoubiquitination assays showed that under our experimental conditions, these two mutants have similar activity as the wild type complex (**Figure 7I and Supp. Figure 4**).

Taken together, these results suggest that disease-relevant mutations in the RYBP ‘N-term helix’ critically destabilize the structure of vPRC1. The Q60H mutant completely prevents complex formation. Other mutants (A63T and Q69K) display less pronounced effects, and no evident loss of enzymatic activity *in vitro*, but do nonetheless induce appreciable structural destabilization.

## DISCUSSION

In this work, we report the topological mapping by XL-MS and SAXS of a vPRC1 isoform (PRC1.4), we determine the conformational propensities of its intrinsically disordered regulatory subunit RYBP by NMR, and we identify novel structural interactions and regulatory structural motifs in RYBP, responsible for the assembly of vPRC1.

Most unexpectedly, we identify novel putatively structured regions in RING1B and RYBP by AlphaFold2. These regions, because of their structural disorder predicted by all other available methods to date, had so far been disregarded in terms of their potential structural and functional role in the context of vPRC1 assembly and more generally of RING1B and RYBP functions. Our study is thus an important representative example of how the novel artificial-intelligence-driven algorithm AlphaFold2 can help guide the characterization of otherwise experimentally very challenging protein complexes. In the last couple of years, AlphaFold2 has already proven to be a crucial tool in structural biology, because the resulting predicted structures can be used as starting models for molecular replacement in X-ray crystallography, for modelling cryo-electron microscopy 3D reconstructions or SAXS 3D volumes, and for the rational design of mutations and functional assays. For example, AlphaFold2 helped deciphering the structure of the human nuclear pore complex (Mosalaganti et al., 2022) or of the full-length gamete surface protein Pfs48/45, a malaria vaccine candidate (Ko et al., 2022). Our study adds to such already remarkable impact, by providing an example of how AlphaFold2 can identify putatively structured regions embedded in regions of high intrinsic disorder. In our case, AlphaFold specifically predicts that the RYBP ‘N-term helix’ motif adopts a helical conformation, but by NMR we experimentally determined that the helical population of this motif is present at very low percentages. This apparent discrepancy between the predictions and experimental data suggests that the RYBP ‘N-term helix’ may be a conditionally folded helix, i.e. that this region adopts a helical structure only within certain environments, i.e. upon PRC1 trimer formation, as recently demonstrated for other similar motifs (Alderson et al., 2023). Considering that about 34% of the human proteome codes for intrinsically disordered regions, with ∼47% of proteins containing long disordered regions (≥30 disordered residues, with an average length of 214 residues, (Alshehri et al., 2020)), our study shows how important it is to systematically characterize those regions computationally and experimentally.

Nonetheless, current AI-modelling tools also display some limitations, namely in predicting the overall structure of protein complexes, as these are unable to predict stoichiometry, alternative homo-oligomeric states and asymmetry. Moreover, the AlphaFold2 results are prediction, with an inherent level of error despite quality scores to estimate model accuracy, which as such require experimental validation. Also in this respect our study is exemplary. Indeed, our work shows that out of the two novel motifs identified by AlphaFold2 in vPRC1.4, so far, only the role and molecular features of the RYBP ‘N-term helix’, but not the RING1B ‘linker helix’ could be validated experimentally.

By integrating biochemical, biophysical and NMR studies with structural mutagenesis, we prove that the RYBP ‘N-term helix’ is essential for vPRC1 complex assembly. This study highlights the importance of NMR as a structural technique suitable for the characterization of intrinsically disordered proteins (IDPs), as NMR can be used to determine the structural propensities and conformational dynamics of IDPs or IDP complexes at atomic resolution, elucidating their interactions mechanisms (e.g. folding-upon binding, highly dynamic complexes, multivalent interactions, regulatory switches triggered by post-translational modifications) and conformational dynamics on multiple time scales (Camacho-Zarco et al., 2022; Schneider et al., 2019). IDPs adopt a vast landscape of conformations intrinsic to their flexibility, that are nonetheless important for their specific biological functions in a vast range of molecular processes, namely in neurodegenerative diseases, molecular recognition, signaling pathways and transcription and replication. An example of this is the role of intrinsic disorder in the transcription and replication machinery of measles virus (Jensen et al., 2011) or in the signaling complex formed between the mitogen-activated protein kinase (MAPK) p38α and the intrinsically disordered N-terminal regulatory domain of the MAPK kinase MKK4 (Delaforge et al., 2018).

Coupled to biochemical and structural studies, our enzymatic characterization of the RYBP ‘N-term helix’ provides further insights into the functional roles that this novel motif may play to regulate vPRC1 function. We have found that one mutant (Q60H) completely prevents complex formation, thus it can be expected that *in vivo* this mutant would inactivate vPRC1. Curiously, this mutant occurs in cancer, suggesting that it is not embryonic lethal, contrary to *rybp* or *ring1b* homozygous null embryos, *rybp* heterozygous mutants (semi-penetrant lethal at birth), or even knockouts of both *ring1b* and *ring1a* (Bajusz et al., 2018). The effects observed upon mutations on vPRC1 subunits may be dependent on the specific sets of genes regulated at different developmental stages and on the role of vPRC1 in lineage commitment, from pluripotent to differentiated cells, as the roles of PRC1 proteins are unique in each stem cell type, equally varying within the same cell type, depending on the developmental stage and environmental stimuli. Other mutants induce a less pronounced structural destabilization, which has negligible catalytic effects *in vitro*, but may nonetheless represent a vulnerability of vPRC1 in the cell and thus contribute to the inefficiency of the complex and the pathogenicity of the resulting mutants. It should also not be excluded that such mutations may down- or up-regulate the RYBP expression levels. Altered RYBP levels would have an impact in global chromatin compaction and thus global alterations of the cellular transcriptome, which are connected to cancers (e.g. liver carcinoma; lung, prostate, breast, cervical cancers; Hodgkin’s lymphoma), and other diseases like chronic rhinosinusitis or type 2 diabetes (Simoes da Silva et al., 2018). These considerations regarding RYBP reflect other similar observations done on other subunits of PRC1. For instance, a recent study has shown that a mutation in the nucleosome-interacting region of RING1A (the paralog of RING1B), in a patient bearing neurological, skeletal, and immunological abnormalities, caused RING1A to preserve its ability to catalyze ubiquitin chain formation, while no longer being able to ubiquitinate H2A in nucleosomes (Pierce et al., 2018).

Finally, besides the novel identification of the RYBP ‘N-term helix’ and its biochemical, structural and functional characterization, our study also provides the first complete vPRC1 inter-subunit interaction map. Our data reveal novel interactions between the PCGF4 RAWUL domain and both RYBP and RING1B Zn-fingers, between the PCGF4 RAWUL domain and the RYBP β-hairpin, between RYBP Zn-finger and the PCGF4 Zn-finger and Pro/Ser-rich regions, regions currently not covered by available crystal structures. At a time when the full assembly of vPRC1 is not known at high-resolution, these data are important to gain an at least preliminary insight into the vPRC1 topology, which will be helpful in the future to guide further functional characterization of the complex and perhaps model a more accurate vPRC1 high-resolution structure. In this context, for instance, the previously-uncharacterized interaction between the PCGF4 RAWUL and the RYBP β-hairpin, mutually exclusive with the RING1B RAWUL/RYBP β-hairpin interaction, is curious and its physiological significance should be investigated more in detail in the future. Indeed, the close structural and sequence similarity across RAWUL domains of various *Polycomb* proteins (Gray et al., 2016; Junco et al., 2013; Wang et al., 2010), coupled to the fact that we identify this interaction not only by AlphaFold2 computational predictions and by NMR titration of individual isolated domains, but also by XL-MS in the context of the fully-assembled vPRC1 trimer, opens up for the possibility of a very high structural and dynamic equilibrium between two conformations of the complex, in which PCGF and RING1B alternatively bind to the same RYBP β-hairpin. Such an exchange may also occur in other vPRC1 isoforms.

In summary, our analysis of the vPRC1.4 isoform provides a topological map of this important chromatin remodeling complex, revealing novel interactions between regions of currently unavailable high-resolution 3D structures, and identifying novel specific structural motifs that are crucial in ensuring complex assembly and thus functionality. These studies help rationalizing the physio-pathological effects of cancer-related mutations in the vPRC1 regulatory subunit RYBP.

## MATERIAL AND METHODS

### Cloning, protein expression and purification

#### vPRC1.4 WT and mutants

cDNA I.M.A.G.E. clones of each human vPRC1.4 subunits were purchased from Source BioScience (Nottingham, UK). The I.M.A.G.E. clone numbers are: RING1B = 4285715, PCGF4 = 4138748 and RYBP = 4813939. The cDNA of each subunit was then cloned into MultiBac system vectors (Bieniossek et al., 2012; Vijayachandran et al., 2011) by sequence and ligation independent cloning (SLIC), yielding pACEBac1-RING1B (acceptor), pIDS-PCGF4 (donor) and pIDK-RYBP (donor) vectors, with RING1B and RYBP constructs containing a TEV protease cleavable N-terminal Strep-tag or 6x-His-tag, respectively. The obtained vectors were then fused *in vitro* by Cre-loxP recombination, as previously described (Haffke et al., 2013), originating a single plasmid containing all three expression cassettes and yielding the formation of vPRC1.4 complex. The generated constructs were transformed into chemically competent DH10 EmBacY cells (harbouring the baculoviral EmBacY genome) (Trowitzsch et al., 2010) and positive clones identified by blue/white screening (in the presence of isopropyl-β-D-thiogalactopyranoside and BluoGal), followed by restriction enzymes digestion and DNA sequencing, to verify for the presence of the genes encoding each subunit of interest and to ensure an in-frame insert. Recombinant baculovirus were produced as previously described (Bieniossek et al., 2012; Trowitzsch et al., 2010) and used to infect Sf21 insect cells at a cell density of 1.0 x 10^6^ cells/mL in Sf-900 SFM medium (Gibco Life Technologies), shaking at 90 rpm on Sartorius Stedim Biotech “CERTOMAT RM” with a 25mm orbit. Protein expression was followed by monitoring YFP (Yellow Fluorescent Protein) expression from the viral backbone and cells harvested between 72h-84h after cell proliferation arrest by centrifugation at 1200g for 15min at 4°C (Beckman Coulter centrifuge, JLA8.1000). Further, vPRC1.4-RYBP single point mutants (Q60H, A63T and Q69K) were generated using the vPRC1.4 recombined plasmid (containing all three expression cassettes) as template and using Phusion polymerase (NEB) according to the manufacturer’s protocol. The oligonucleotides used for mutagenesis are provided in **Table 1**. The generated mutants were expressed as described above for the wild type vPRC1.4 complex.

**Table 1:**
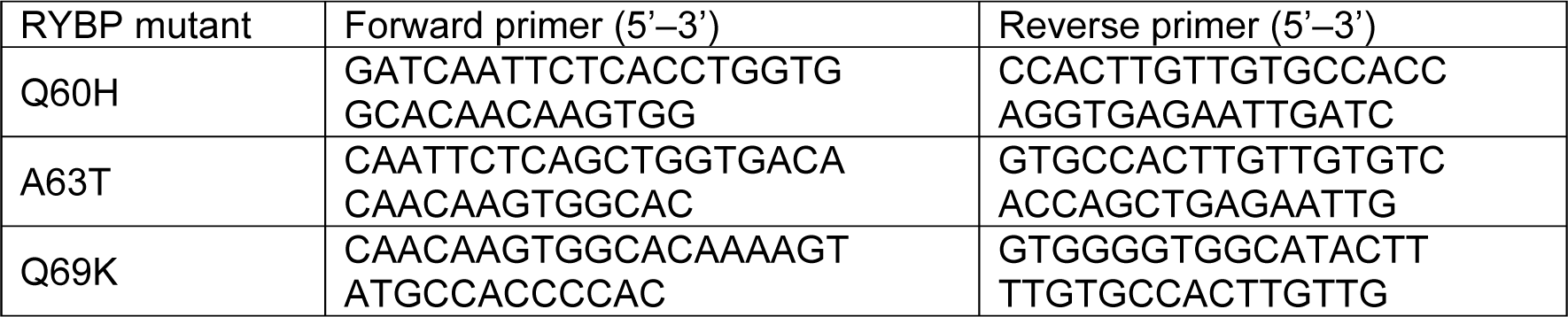
Primers used to produce vPRC1-RYBP mutants used in this study.

vPRC1.4 pellets from 800-1000 x 10^6^ harvested cells were resuspended in 100mL of lysis buffer (50mM Tris pH 9.0, 150mM NaCl, 1mM DTT supplemented with complete EDTA-free protease inhibitors cocktail and DNAseI). Cells were disrupted by sonication on ice, for 3min and 30s at 30% amplitude, using a Sonics VCX-750 Vibra Cell Sonicator (Sonics & Materials Inc., Newtown, CT, USA), followed by centrifugation at ∼28500g for 1h at 4°C (Beckman Coulter centrifuge, JA-14) in order to remove cell debris. The supernatant was incubated for 1h, stirring at 4°C, with 3mL of Ni-Sepharose affinity resin beads pre-equilibrated with 50mM Tris pH 9.0, 150mM NaCl, 1mM DTT (buffer A), and the mix then loaded onto a BioRad gravity flow column and the respective flow-through collected. The resin was washed with 3CV of buffer A, followed by ∼7CV of wash buffer (buffer A with 20mM imidazole). The protein complex was subsequently eluted, pulled by RYBP subunit, with 10CV of elution buffer (buffer A containing 100mM imidazole). A final wash step of ∼7CV with 50mM Tris pH 9.0, 150mM NaCl, 1mM DTT, 400mM imidazole was applied to the resin in order to deplete it of any bound protein. The elution fraction was then concentrated using an amicon of 30 kDa MWCO (Eppendorf 5804R, ∼3000g) at 4°C, before being loaded onto a size exclusion Superdex 200 HiLoad 16/600pg column (GE-Healthcare), pre-equilibrated with 50mM Tris pH 9.0, 150mM NaCl. Final fractions of interest containing vPRC1.4 were pooled, concentrated and used for subsequent experiments. vPRC1.4-RYBP single point mutants were all isolated following the same purification protocol.

#### Production of ^15^N- or ^13^C/^15^N-labelled RYBP

RYBP full-length coding sequence was cloned into the *Escherichia coli* (*E.coli*) expression vector pESPRIT002 (Hart and Tarendeau, 2006), using the AatII and NotI restriction sites. The plasmid contains an N-terminal 6x-His-tag followed by a TEV protease cleavage site. RYBP FL subunit was used for NMR experiments and expressed in M9 medium, either ^15^N- or ^13^C/^15^N-labelled, using the bacterial strain BL21 (DE3) pLysS cells.

Cells expressing RYBP were grown in M9 medium (10 g/L Na_2_HPO_4_.7H_2_O, 3 g/L KH_2_PO_4_, 0.5 g/L NaCl, 1 g/L ^15^NH_4_Cl) supplemented with 2 g/L of D-glucose or ^13^C D-glucose, 1mM MgSO_4_, 0.1mM CaCl_2_, 0.1mM MnCl_2_, 50µM ZnSO_4_, 100µM FeCl_3_, vitamins cocktail (M9 enriched medium) and antibiotics (chloramphenicol 34 µg/mL and kanamycin 50 µg/mL). An over-day LB preculture containing chloramphenicol and kanamycin was launched and left growing for 8h at 37°C, 160rpm. Cells were spun down for 15min at 3000g (Eppendorf 5804R) and inoculated in M9-^15^N or M9-^13^C/^15^N enriched medium with antibiotics and left growing overnight at 37°C, 160rpm. The following day, overnight precultures were inoculated in culture flasks containing M9-^15^N or M9-^13^C/^15^N enriched medium (1:20) with antibiotics, and cells grown at 37°C, 140rpm, until an OD_600_ of 0.6-0.8 was reached. The temperature was then reduced to 20°C and protein expression induced by addition of 0.5mM isopropyl-β-D-thiogalactopyranoside (IPTG). Expression was continued for 7h and the cells harvested by centrifugation at ∼6250g for 15 min at 4°C (Beckman Coulter centrifuge, JLA8.1000).

Harvested cells were resuspended in lysis buffer (50mM Tris pH 9.0, 150mM NaCl, 1mM DTT, supplemented with complete EDTA-free protease inhibitors cocktail, DNAseI, 1mM leupeptin and 1mM pepstatin) and the cells lysed using a microfluidizer (9000 psi process pressure; Microfluidics M-110L microfluidizer), after three passage rounds. The lysate was then centrifuged at ∼28500g for 1h at 4°C (Beckman Coulter centrifuge, JA-14) and the supernatant incubated for 1h, stirring at 4°C, with 14mL of Ni-Sepharose affinity resin beads pre-equilibrated with buffer A (50mM Tris pH 9.0, 150mM NaCl, 1mM DTT, complete EDTA-free protease inhibitors cocktail, 1mM leupeptin and 1mM pepstatin). The sample/resin mix was applied onto a BioRad glass Econo-column and the flow-through collected. The resin was further washed with 2CV of buffer A, followed by 4CV of wash buffer (buffer A with 20mM imidazole), and RYBP subsequently eluted with 6.5CV of elution buffer (buffer A containing 200mM imidazole). A final wash step of ∼2.5CV with buffer A containing 400mM imidazole was applied to the resin in order to deplete it of any remaining protein. Protein fractions of interest were pooled and further purified through a 5mL heparin column (GE-Healthcare) pre-equilibrated with buffer A, and RYBP eluted against a 7CV linear salt gradient, spanning from 150mM to 1M NaCl, followed by a gradient delay of 3CV at 1M NaCl. RYBP fractions of interest were pooled after the heparin column and concentrated using an amicon of 10 kDa MWCO (Eppendorf 5804R, ∼3000g) at 4°C, before being loaded onto a size exclusion Superdex 200 HiLoad 16/600pg column (GE-Healthcare), pre-equilibrated with 50mM Tris pH 6.5, 150mM NaCl, complete EDTA-free protease inhibitors cocktail, 1mM leupeptin and 1mM pepstatin, as a final purification step. Final fractions of interest containing RYBP were pooled, concentrated and used for subsequent NMR experiments, after addition of 90% H_2_O/10% D_2_O.

#### RING1B-PCGF4 dimer

A subcomplex of vPRC1.4, containing the subunits RING1B and PCGF4 (RING1B-PCGF4 dimer) was generated following the same cloning strategy and expression protocols as for the vPRC1.4 complex. Although known to be successfully purified via a two-step purification protocol involving a strep-tactin affinity column followed by size exclusion chromatography, as previously described (Colombo et al., 2019), such protocol does not produce enough yield for proceeding to NMR experiments. Hence, for such subcomplex, insect-cell-expressed RING1B-PCGF4 dimer was co-purified with *E.coli* expressed RYBP (^15^N- or ^13^C/^15^N-labelled), following the same purification protocol as the one above described for RYBP subunit alone. Final fractions containing the proteins of interest were pooled, concentrated and used for subsequent NMR experiments.

#### RING1B and PCGF4 subunits subdomains

The RAWUL domain of both RING1B and PCGF4 subunits were also cloned into the expression vector pESPRIT002, following the same strategy as for RYBP, with both constructs containing a TEV protease cleavable N-terminal 6x-His-tag. RING1B RAWUL [223-333]a.a. and PCGF4 RAWUL [121-235]a.a. domain containing constructs were overexpressed in *E.coli* BL21 (DE3) pLysS cells. Cells were grown at 37°C, 140rpm, in Luria-Bertani (LB) or Terrific Broth (TB) culture medium, for RING1B and PCGF4 respectively, supplemented with chloramphenicol (34 µg/mL) and kanamycin (50 µg/mL), until an OD_600_ of 0.7-0.8 for LB, or 1.0-1.5 for TB medium. The temperature was then reduced to 18°C and protein expression induced by addition of 0.5mM IPTG. Expression was continued for 16h and the cells harvested by centrifugation at ∼6250g for 15 min at 4°C (Beckman Coulter centrifuge, JLA8.1000). The RING Zn-finger domains from both RING1B and PCGF4 subunits were cloned into a modified polycistronic expression vector (pET29a) by restriction free cloning using NEB Builder HiFi DNA Assembly kit, and inserting a TEV protease cleavable 6x-His-tag at the N-terminus of RING1B Zn-finger. The resulting construct, co-expressing RING1B Zn-finger [14-116]a.a. and PCGF4 Zn-finger [1-103]a.a., was overexpressed in BL21 (DE3) pLysS cells. Cells were grown at 37°C, 140rpm, in LB medium with chloramphenicol and kanamycin and until an OD_600_ of 0.7-0.8. Protein induction was obtained by adding 0.5mM IPTG and overnight culture growth at 37°C, for further 16h. Cells were harvested by centrifugation at ∼6250g for 15 min at 4°C (Beckman Coulter centrifuge, JLA8.1000).

Cells expressing RING1B RAWUL [223-333]a.a. domain were resuspended in lysis buffer (50mM Tris pH 8.0, 150mM NaCl, 1mM DTT, supplemented with complete EDTA-free protease inhibitors cocktail and DNAseI) and lysed using a microfluidizer (9000 psi process pressure; Microfluidics M-110L microfluidizer), with three passage rounds. After lysis, the lysate was centrifuged at ∼18000g for 1h at 4°C (Beckman Coulter centrifuge, JA-20) and the supernatant incubated for 45min, stirring at 4°C, with 3mL of Ni-Sepharose affinity resin beads pre-equilibrated with buffer A (50mM Tris pH 8.0, 150mM NaCl, 1mM DTT). Supernatant/resin mix was then loaded onto a BioRad gravity flow column and the respective flow-through collected. The resin was further washed with 7CV of buffer A, followed by ∼13CV of wash buffer (buffer A with 5mM imidazole). Protein was then eluted with 12CV of elution buffer (buffer A with 100mM imidazole) and a final wash step of 5CV, with buffer A containing 400mM imidazole, was applied to the resin in order to deplete it of any remaining protein. Protein fractions of interest were pooled and incubated overnight at 4°C with 2% (w/w) 6x-His-tagged TEV protease, dialysing against buffer A and using a 6-8 kDa cut-off dialysis membrane, in order to cleave the N-terminal 6x-His-tag. The untagged protein was recovered in the flow-through, after a reverse Ni-Sepharose column, and concentrated using an amicon of 10 kDa MWCO (Eppendorf 5804R, ∼3000g) at 4°C, before being loaded into a size exclusion Superdex 75 HiLoad 16/600pg column (GE-Healthcare), pre-equilibrated with 50mM Tris pH 6.5, 150mM NaCl. Final fractions containing RING1B RAWUL [223-333]a.a. domain were pooled, concentrated and used for subsequent NMR experiments.

Pellet resuspension and lysis of cells expressing PCGF4 RAWUL [121-235]a.a. domain were done as for RING1B RAWUL domain. After lysis, the lysate was centrifuged at ∼28500g for 1h30 at 4°C (Beckman Coulter centrifuge, JA-14) and the supernatant incubated for 1h, stirring at 4°C, with 8mL of Ni-Sepharose affinity resin beads pre-equilibrated with buffer A (50mM Tris pH 8.0, 150mM NaCl, 1mM DTT). Supernatant/resin mix was then loaded onto a BioRad glass Econo-column and the respective flow-through collected. The resin was further washed with 5CV of buffer A, followed by two incremental wash steps, one with 10mM imidazole, and the second one with 20mM imidazole, both for ∼12CV. Protein was eluted with 9CV of elution buffer (buffer A with 250mM imidazole) and a final wash step of 5CV, with buffer A containing 400mM imidazole, was applied to the resin in order to deplete it of any remaining protein. In order to cleave PCGF4 RAWUL N-terminal 6x-His-tag, fractions of interest were pooled and dialysed overnight at 4°C, against dialysis buffer (50mM Tris pH 8.0, 75mM NaCl, 1mM DTT) and in the presence of 3% (w/w) 6x-His-tagged TEV protease, using a 6-8 kDa cut-off dialysis membrane. Additional 2h of cleavage were performed at room temperature the next day. Protein sample was then further purified through a 5mL HiTrap SP HP cation exchange column (GE-Healthcare) pre-equilibrated with 50mM Tris pH 8.0, 75mM NaCl. PCGF4 RAWUL domain eluted against a 7CV linear salt gradient, spanning from 75mM to 1M NaCl, followed by a gradient delay of 3CV at 1M NaCl. Protein fractions of interest were pooled and concentrated using an amicon of 10 kDa MWCO (Eppendorf 5804R, ∼3000g) at 4°C, before being loaded onto a size exclusion Superdex 75 10/300GL column (GE-Healthcare), pre-equilibrated with 50mM Tris pH 6.5, 150mM NaCl. Final fractions containing PCGF4 RAWUL [121-235]a.a. domain were pooled, concentrated and used for subsequent NMR experiments.

As for RING1B and PCGF4 RAWUL domains, cells expressing RING1B/PCGF4 Zn-finger dimer were resuspended in lysis buffer (50mM Tris pH 8.0, 150mM NaCl, 1mM DTT, supplemented with complete EDTA-free protease inhibitors cocktail and DNAseI). Cells were lysed using a microfluidizer (9000 psi process pressure), followed by lysate centrifugation at ∼28500g for 1h at 4°C (Beckman Coulter centrifuge, JA-14). The supernatant was incubated for 45min, stirring at 4°C, with 4mL of Ni-Sepharose affinity resin beads pre-equilibrated with buffer A (50mM Tris pH 8.0, 150mM NaCl, 1mM DTT), and the mix then loaded onto a BioRad gravity column and the respective flow-through collected. The resin was further washed with ∼3CV of buffer A, followed by two incremental wash steps, one with 30mM imidazole, and the second one with 50mM imidazole, both for 10CV. Protein was eluted with ∼13CV of elution buffer (buffer A with 200mM imidazole) and a final wash step of 4CV, with buffer A containing 400mM imidazole, was applied to the resin in order to deplete it of any remaining protein. Protein fractions of interest were pooled and incubated overnight at 4°C with 3% (w/w) 6x-His-tagged TEV protease, dialysing against 50mM Tris pH 8.0, 100mM NaCl, 1mM DTT and using a 6-8 kDa cut-off dialysis membrane, in order to cleave the N-terminal 6x-His-tag. Additional 2h of cleavage were performed at room temperature the next day. Protein sample was then further purified through a 5mL HiTrap SP HP cation exchange column (GE-Healthcare), following the same protocol as for PCGF4 RAWUL domain, with a linear salt gradient spanning from 100mM to 1M NaCl. Protein fractions of interest were pooled and concentrated using an amicon of 10 kDa MWCO (Eppendorf 5804R, ∼3000g), before being loaded onto a size exclusion Superdex 75 HiLoad 16/600pg column (GE-Healthcare), pre-equilibrated with 50mM Tris pH 6.5, 150mM NaCl. Final fractions containing RING1B/PCGF4 Zn-finger dimer were pooled, concentrated and used for subsequent NMR experiments.

### Analytical Size Exclusion Chromatography and Multi-angle Laser Light Scattering (SEC-MALLS)

Multi-angle laser light scattering (MALLS) coupled with size exclusion chromatography (SEC) and measurement of refractive index (RI) allows the determination of the absolute molecular mass of a particle in solution, independently of its dimensions and shape. Size exclusion chromatography was performed on a high-pressure liquid chromatography system equipped with an LC-20AD pump, an autosampler SIL20-AC_HT_ storing the samples at 4°C, a communication interface CBM20A (Shimadzu), using a Superdex 200 10/300GL column (GE Healthcare) pre-equilibrated with 50mM Tris pH 9.0, 150mM NaCl and thermostated at 20°C in an oven XLTherm (WynSep). Analytical runs were performed at 20°C with a flowrate of 0.5mL/min. Typically, 50µL of vPRC1.4 complex, or vPRC1.4-RYBP point mutants, at a concentration between 1.8-2.5 mg/mL was injected. The elution profiles were followed on-line at 280 nm (SPD-M20A Shimadzu) with both static and dynamic light scattering being detected by a miniDAWN-TREOS and a Dynapro Nanostar detector (Wyatt Technology Corp., Santa Barbara, CA), respectively, using a laser emitting at 658 nm. Protein concentration was measured on-line with the use of differential refractive-index measurements, with an Optilab rEX detector (Wyatt Technology Corp., Santa Barbara, CA) and an RI increment, dn/dc, of 0.187 mL/g. Data were analysed and weight-averaged molar masses (Mw) calculated using ASTRA software (Wyatt Technology Corp., Santa Barbara, CA).

### Native Mass Spectrometry (Native MS)

vPRC1.4 complex was analysed by native MS at a concentration of 4µM. Protein buffer was freshly exchanged by gel filtration to 250mM ammonium acetate with 1mM DTT. Protein ions were generated using a nanoflow electrospray (nano-ESI) source. Nanoflow platinum-coated borosilicate electrospray capillaries were bought from Thermo Electron SAS (Courtaboeuf, France). MS analyses were carried out on a quadrupole time-of-flight mass spectrometer (Q-TOF Ultima, Waters Corporation, Manchester, U.K.). The instrument was modified for the detection of high masses (Boeri Erba et al., 2018; Boeri Erba et al., 2020; Sobott et al., 2002; van den Heuvel et al., 2006). The following instrumental parameters were used: capillary voltage = 1.2–1.3 kV, cone potential = 40 V, RF lens-1 potential = 40 V, RF lens-2 potential = 1 V, aperture-1 potential = 0 V, collision energy = 30–140 V, and microchannel plate (MCP) = 1900 V. All mass spectra were calibrated externally using a solution of cesium iodide (6 mg/mL in 50% isopropanol) and were processed with the Masslynx 4.0 software (Waters Corporation, Manchester, U.K.) and with Massign software package (Morgner and Robinson, 2012).

### Analytical Ultracentrifugation (AUC)

Purified vPRC1.4 sample was subjected to analytical ultracentrifugation experiments at a protein concentration of 1mg/mL. Sedimentation-velocity experiments were performed overnight at a rotor speed of 42000rpm and 4°C, using an analytical ultracentrifuge XL-I with an Anti-60 rotor (Beckman Coulter) and double-sector cells (Nanolytics) of optical path length 12mm with Saphire windows. Data acquisitions were made using absorbance at 280nm and interference optics. A partial specific volume (ῡ) of 0.718 mL/g, buffer density (ρ) of 1.01 g/mL and viscosity (η) of 1.61 cp were calculated from the sample composition with the programs Sedfit and Sednterp. Data were analysed using Sedfit software, using continuous c(s) distribution model of sedimentation coefficients (s) (Schuck, 2000).

### Size exclusion chromatography coupled to small-angle X-ray scattering (SEC-SAXS)

Size exclusion chromatography coupled with small-angle X-ray scattering experiments were conducted on the BioSAXS BM29 beamline at the European Synchrotron Radiation Facility (ESRF, Grenoble, France). An on-line HPLC system was attached directly to the sample inlet valve of the beamline sample changer. Purified vPRC1.4 protein sample (50µL) at 1mg/mL, previously spun down (Eppendorf 5427R, FA-45-30-11, 18000*g*, 10min), was manually injected onto a Superdex 200 increase 5/150 GL column (GE Healthcare), pre-equilibrated with 50mM Tris pH 9.0, 150mM NaCl, 1mM DTT buffer. Buffers were degassed prior to the run and a flow rate of 0.2mL/min at 4°C was used. The baseline was monitored prior to the experimental run. All data from the run were collected at a wavelength λ=0.99 Å, using a sample-to-detector (PILATUS 1M; Dectris AG, Baden, Switzerland) distance of 2.87m corresponding to a q-range of 0.0035-0.167 Å^-1^ where q is the momentum transfer (q= 4πλ sinθ) and 2θ the scattering angle. Acquisitions with an exposure time of 1s/frame were recorded during 1000s. One hundred initial frames were averaged to create the reference buffer and the frames collected at the elution peak, corresponding to the scattering of an individual purified species, were also averaged (to give the average scattering curve) and subtracted of the reference buffer using the program PRIMUS (Konarev et al., 2003). Primary data reduction was performed and Radii of gyration (R_g_) and pairwise distance distribution functions [P(r)] (D_max_) were extracted based on the Guinier approximation.

### Crosslinking mass spectrometry (XL-MS)

About 50µg of purified vPRC1.4 complex was crosslinked by addition of 5µL at 50mM of an iso-stoichiometric mixture of H12/D12 isotope-coded di-succinimidyl-suberate (DSS) and incubated for 30min at 37°C. Reaction was quenched by addition of ammonium bicarbonate (NH_4_HCO_3_) to a final concentration of 50mM for 10 min at 37°C. Crosslinked proteins were denatured using urea and Rapigest at a final concentration of 4 M and 0.05% (w/v), respectively. Samples were reduced with 10 mM DTT for 30 min at 37°C, followed by cysteines carbamidomethylation with 15 mM iodoacetamide treatment for 30 min at room temperature in the dark. Protein digestion was then performed, first using 1:100 (w/w) LysC (Wako Chemicals, Neuss, Germany) for 4 h at 37°C, and finalised with 1:50 (w/w) trypsin digestion overnight at 37°C, after the urea concentration was diluted to 1.5 M. Samples were then acidified with 10% (v/v) trifluoroacetic acid (TFA) and desalted using OASIS® HLB µElution Plate. Crosslinked peptides were desalted and reconstituted with SEC buffer [30% (v/v) ACN in 0.1% (v/v) TFA] and fractionated using a Superdex Peptide PC 3.2/30 column (GE Healthcare). Collected fractions were analysed by liquid chromatography-coupled tandem mass spectrometry (MS/MS) using a nanoAcquity UPLC system connected on-line to LTQ-Orbitrap Velos Pro instrument. To assign the fragment ion spectra, raw files were converted to centroid mzXML using a raw converter and then searched using xQuest (Leitner et al., 2014) against a FASTA database containing the sequences of the crosslinked proteins. Posterior probabilities were calculated using xProphet, and results were filtered using the following parameters: FDR = 0.05, min delta score = 0.95, MS1 tolerance window of 4–7 p.p.m., ld score > 25.

### Limited Proteolysis

The vPRC1.4 complex (1 mg/ml) was treated with trypsin at an enzyme-to-protein ratio of 1:10 (w/w). Samples were taken after 2, 5, 10, 20, 30, 40, 60 and 120 min and analyzed by SDS–PAGE. The proteolyzed samples were extracted from the gel and analyzed by PMF-MS at the EMBL Proteomics Facility (EMBL Heidelberg).

### Evolutionary Sequence Alignment Analysis

Sequence alignments were performed with Clustal Omega (Madeira et al., 2022) using the default parameters for multiple alignment sections. Protein sequences used were obtained from GenBank. Resulting alignments and secondary structure predictions were rendered with ESPript3 (Robert and Gouet, 2014). The sequences used for RYBP sequence alignments are as follows: Human (*Homo sapiens*, AAH36459.1), Gorilla (*Gorilla gorilla*, XP_030865299.1), Orangutan (*Pongo abelii*, XP_024100611.1), Mouse (*Mus musculus*, XP_036014155.1), Rat (*Rattus norvegicus*, NP_001386529.1), Sheep (*Ovis aries*, XP_027813966.1), Opossum (*Monodelphis domestica*, XP_056660407.1), Chicken (*Gallus gallus*, XP_046755973.1), Clawed Frog (*Xenopus laevis*, XP_018116960.1), Zebrafish (*Danio rerio*, NP_001012373.1), Platy fish (*Xiphophorus maculatus*, XP_023180825.1) and Placozoa (*Trichoplax adhaerens*, XP_002111440.1). For RING1B sequence alignments the sequences used are: Human (*Homo sapiens*, NP_009143.1), Gorilla (*Gorilla gorilla*, XP_030859631.1), Orangutan (*Pongo abelii*, NP_001127433.1), Mouse (*Mus musculus*, NP_001347773.2), Rat (*Rattus norvegicus*, NP_001388309.1), Sheep (*Ovis aries*, XP_027831758.1), Opossum (*Monodelphis domestica*, XP_001366901.1), Chicken (*Gallus gallus*, XP_015146036.1), Clawed Frog (*Xenopus laevis*, XP_018116603.1), Zebrafish (*Danio rerio*, NP_571288.2), Platy fish (*Xiphophorus maculatus*, XP_005809264.1) and Placozoa (*Trichoplax adhaerens*, XP_002108615.1). Numbers indicated correspond to GenBank accession numbers.

### Nucleosome production

About 50 µL of TOP10 competent cells were transformed with 10 ng of pST55 plasmid containing 16 copies of 147 bp 601 Widom DNA sequence (Lowary and Widom, 1998). Widom 601 DNA was then purified as previously described (McGinty et al., 2016). Nucleosomes (NCP) were reconstituted as described (Shim et al., 2012).

### H2A monoubiquitination activity assay

A 3x-concentrated master mix containing 50mM Na-HEPES pH 7.7, 90 nM E1 enzyme (BML-UW9410-0050; Enzo Life Sciences, Villeurbanne, France), 1.2 µM of E2 enzyme UbcH5c, 10 mM DTT, 6 mM ATP, 30 µM ZnSO4, 15 mM MgCl_2_, 23 µM methylated ubiquitin (U-501-01M; R&D System, Minneapolis, MN, USA) was pre-heated at 37°C for 20 min. vPRC1.4 WT, vPRC1.4-RYBP point mutants and reconstituted vPRC1.4 were prepared at 10 concentrations (0.6, 0.9, 1.2, 1.5, 1.8, 2.1, 2.4, 3.0, 3.6, 4.2 µM) in final buffer 50mM Tris pH 9.0, 150mM NaCl. Nucleosomes were prepared at 2.1 µM, buffer exchanged into 50mM Tris pH 9.0, 150mM NaCl, 0.5M TCEP. Finally, 5 µL of master mix 3x, 5 µL of each vPRC1.4 complex prepared concentrations and 5 µL of nucleosomes were mixed and incubated at 37°C for 100 min. The reaction was quenched by adding SDS buffer (5% glycerol, 2% β-mercaptoethanol, 50 mM Tris-HCl pH 8.0, bromophenol blue, 2% SDS) and boiling at 95°C. All samples were loaded on pre-casted SDS Tris-glycine 4–20% polyacrylamide gels (Life Technologies, Carlsbad, CA, USA). For western blot detection, gels were transferred onto nitrocellulose membranes (Trans-Blot Turbo, Mini-Size Nitrocellulose, Bio-Rad) using Trans-Blot Turbo Mini-size Transfer stacks (Bio-Rad) and the Trans-Blot Turbo Transfer system (Bio-Rad), for 7min (1.3A constant, up to 25V). Each membrane was blocked overnight with a 2% solution of BSA diluted in Tris buffer saline with Tween 20 at 0.1% (TBST) at 4°C. The membranes were washed 3x for 8min with TBST and incubated with the primary antibody mix containing anti-H2AUb antibody (1:600, 06-678; Millipore, Burlington, MA, USA) and anti-H2A (1:2500, 07-146; Sigma Aldrich, Saint-Louis, MO, USA) for 4h at room temperature. The anti-H2A antibody displays cross-reactivity with histone H4 and is outcompeted by the anti-H2AUb antibody, when used simultaneously on ubiquitinated H2A (Rose et al., 2016). Membranes were washed 3x for 8min with TBST and incubated for 2h, at room temperature, with the secondary antibodies anti-mouse (1:5000, A11002; Thermo Fisher Scientific, for anti-H2AUb) conjugated with Alexafluor dye 532 nm and anti-rabbit (1:5000, A32731; Thermofisher, for anti-H2A) conjugated with Alexafluor dye 488 nm. The membranes were washed 3x for 8min with TBST and the fluorescent signal from the membranes was recorded with an Amersham Typhoon (GE Healthcare). For each sample, Mono-Ub activity was measured three times independently. The bands were quantified with ImageQuant (GE Healthcare). The H2A monoubiquitination activity was calculated as the ratio between ubiquitinated and total H2A and plotted over the concentrations of vPRC1.4 complexes. Data were analysed using GRAPHPAD (GraphPad Software).

### Electrophoretic Mobility Shift Assays (EMSA)

Native polyacrylamide gels were used to test interaction between vPRC1.4 complex and nucleosomes (NCP). Nucleosomes concentration was kept constant, while vPRC1.4 concentrations were varied, at increasing vPRC1.4:NCP ratios (1.5, 2.5, 3.5, 5.0, 7.5, 10). vPRC1.4 and NCP binding reactions were incubated on ice for 30min, in 50mM Tris pH9.0, 150mM NaCl, and NCP:vPRC1.4 complexes run on a 6% polyacrylamide gel using TBE buffer 0.25x in non-denaturing conditions, for 1h at 120V and 4°C. Gels were stained with Sybr Safe and scanned with a Geldoc.

### Nuclear Magnetic Resonance (NMR)

NMR spectra were recorded of free RYBP at 5°C and a ^1^H frequency of 950 MHz. The NMR spectral assignment was carried out on a ^13^C/^15^N-labeled sample (concentration of 293 μM) in 50 mM Tris pH 6.5, 150 mM NaCl using a series of BEST-TROSY triple resonance experiments (Solyom et al., 2013): HNCO, intra-residue HN(CA)CO, HN(COCA)CB and intra-residue HN(CA)CB. The assignment was done manually and included all residues of RYBP, except those belonging to the Zn-finger domain for which magnetization transfer was incomplete in the triple resonance experiments. The secondary structure propensities of RYBP were calculated from the experimental ^13^Cα and ^13^Cβ chemical shifts using the SSP program (Marsh et al., 2006).

The ^1^H-^15^N HSQC spectrum of the free RYBP was compared to that of RYBP in the context of the trimeric complex with the RING1B/PCGF4 catalytic dimer of vPRC1.4 (concentration of 203 μM). This comparison allows to identify the regions of RYBP that experience a restriction of motion in the context of the trimeric complex. In addition, we measured ^15^N *R*_1ρ_ relaxation rates in RYBP at 5°C and at a ^1^H frequency of 950 MHz (both in its free form and the context of the trimeric complex) using published ^1^H-^15^N HSQC-based pulse sequences and a spin lock field of 1.5 kHz (Lakomek et al., 2012). The decay of magnetization was sampled at: 1, 10, 30, 50, 70, 90, 130, 170, 200 and 230 milliseconds (ms). A technical replicate was acquired at 70 ms for error estimation.

To further map the interactions within the complex, we carried out titrations of ^15^N-labeled RYBP (50 μM) with increasing concentrations (25, 50 and 100% molar ratios) of RING1 RAWUL [223-333]a.a., PCGF4 RAWUL [121-235]a.a. and RING1B [14-116]a.a./PCGF4 [1-103]a.a. Zn-finger dimer domains. For each titration step, an ^1^H-^15^N HSQC spectrum was recorded. The data were analyzed in terms of intensity ratios (*I*/*I*^0^), where *I* is the intensity in the spectra with added RING1 RAWUL, PCGF4 RAWUL or the Zn-finger dimer, while *I*^0^ is the intensity in the spectrum of free RYBP.

### AlphaFold2

To compute the vPRC1.4 complex structure, using the machine-learning algorithm AlphaFold2, the amino acid sequences of RING1B, PCGF4 and RYBP full-length subunits were submitted to the AlphaFold2 colab notebook (Mirdita et al., 2022) and the 5 resulting models were compared based on their plDDT (predicted local distance difference test) and PAE (predicted alignment error). The prediction failed to identify a confident complex structure. Additionally, changing the order of the protein sequences submitted yielded different results. Finally, we have used AlphaFold2 to predict model structures of each vPRC1.4 subunit in isolation, which allowed us to detect new structural motifs. (https://colab.research.google.com/github/sokrypton/ColabFold/blob/main/AlphaFold2.ipynb#scrollTo=G4yBrceuFbf3)

### Mass Photometry

Mass photometry measurements were performed for vPRC1.4 WT and vPRC1.4-RYBP point mutants samples using a One^MP^ mass photometer (Refeyn Ltd). Immediately prior to mass photometry measurements, protein stocks were diluted to concentrations between 15-20nM (adjusted to maximize camera counts while avoiding saturation) in filtered 50mM Tris pH 9.0, 150mM NaCl buffer, into a final 20μL droplet placed on the gasket. Data acquisition was started within 10s after addition of the protein. Each sample was measured three times independently with similar results. Contrast-to-Mass (C2M) calibration was performed under the same buffer conditions as our samples and using a Native Marker (NM) composed of 4 proteins with molecular weights of 66, 146, 480 and 1048 kDa (Native Mark unstained protein standard, LC0725, Life Technologies). Ratiometric movies were collected for the buffer, native marker and the measured samples for 1min. All measurements were performed using the regular field of view (18μm^2^ of detection area). Data acquisition was performed using AcquireMP (Refeyn) and images processed and analysed using DiscoverMP (Refeyn). Mass Photometry technique has a detection size limit range of 40 kDa to 5 MDa.

### Thermal Shift Assay (TSA)

To compare the thermal stability of the vPRC1.4 wild type and mutants, thermal shift assays were performed using the fluorescent probe SYPRO Orange (Boivin et al., 2013). vPRC1.4 samples were used at final concentrations between 5-10µM in 50mM Tris pH 9.0, 150mM NaCl, to which 5X SYPRO Orange dye (Invitrogen) was added, in a final volume of 20 µL. The thermal stability was measured using a real time PCR machine (CFX96, C1000 Touch, BioRad). The probe was excited at 488 nm and the emission signal was recorded at 585 nm while the temperature was increased by increments of 1°C/min from 25 to 95°C. Each sample was measured at least four times independently. Fluorescence data of each well were corrected for background signal and normalized between 0 and 1. Subsequently, fluorescence values were plotted in function of their corresponding temperature values to visualize thermal shift profile curves. Using the *drc* package (version 3.0.1; (Xia et al., 2015)) in R studio (version 2022.02.0; (Team, 2013)), a five-parameter log-logistic model was fitted on the data for both transitions. The model is based on the following function:

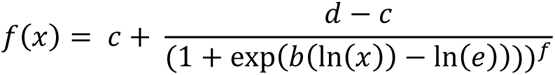

where *b* is the slope; *c* the minimum fluorescent value; *d* the maximum fluorescent value; *e* the inflection point; and *f* the asymmetry factor. Because the inflection point of an asymmetrical sigmoidal curve (*f* ≠ 1) does not coincide with its midpoint, midpoint T_m_ were calculated using the estimated parameters according to the equation:

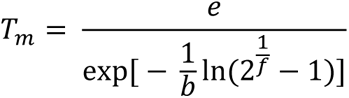

The amount of protein denaturing under each melting transition is proportional to the surface area under the first derivative curve of the fitted function in the interval of T_m_ ± 5°C for the corresponding transition. Therefore, the percentage of protein denaturing at each melting transition was calculated by quantifying the relative proportion of the surface areas of each transition. Standard error of the mean (SEM) of biological replicates was calculated by including one technical replica per biological preparation (n=4). To assess if variances between the mutants were equal, the Levene test, implemented in the statistical software package JMP (JMP Pro version 16.0.0, SAS Institute), was used. Variance between technical replicas of the mutants was calculated by taking the average of the SEM of technical replicas (n=3) of each biological replicate.

## AUTHOR CONTRIBUTIONS

MM, CSS and IGF designed the experiments, CSS, IGF, OP, and ID cloned, expressed and purified all samples and characterized them biochemically, biophysically and enzymatically. LMP and MRJ performed the NMR experiments. EBE performed the native MS experiments. MM supervised the research. CSS and MM wrote the initial draft of the manuscript. All authors approved the final version of the manuscript.

## ACKNOWLEDGEMENTS

We thank all members of the Marcia lab for helpful discussion, the EMBL Proteomic core facility in Heidelberg for support with crosslink- and PMF-MS, Alice Aubert and Dr. Martin Pelosse for support in using the Eukaryotic Expression Facility at EMBL Grenoble, Aline Leroy and Dr. Christine Ebel (IBS Grenoble) for support with AUC, Anna Katharina Hoefler for performing initial evolutionary alignments; Sabina Colombo for help with vPRC1 purification; beamline scientists at the ESRF BioSAXS beamline BM29 for support with SAXS data collection, and Prof. Song Tan (Penn State University) for sharing the pST55 plasmid for nucleosome production. Work in the Marcia lab is partly funded by ITMO Cancer (18CN047-00), Région Auvergne Rhône Alpes (project R21105CC; allocation RPH21004CCA), FINOVI (AAP15), Canceropole CLARA (Oncostarter), and by the Fondation ARC pour la recherche sur le cancer (PJA-20191209284). Financial support from the IR INFRANALYTICS FR2054 for conducting the research is gratefully acknowledged. M.R.J. is laureate of the Impulscience^®^ program of the Fondation Bettencourt Schueller. This work used the platforms of the Grenoble Instruct Center (ISBG: UAR 3518 CNRS-CEA-UJF-EMBL) with support from FRISBI (ANR-10-INBS-0005-02) and GRAL (ANR-10-LABX-49-01) within the Grenoble Partnership for Structural Biology (PSB). The Institute of Structural Biology acknowledges integration into the Interdisciplinary Research Institute of Grenoble.

## Abbreviations

PRC1: *Polycomb* Repressive complex 1
cPRC1: canonical PRC1
vPRC1: variant PRC1
PRC2: *Polycomb* Repressive complex 2
XL-MS: cross linking mass spectrometry
H3K27me3: trimethylated histone H3 on Lys27
H2AK119ub: monoubiquitinated H2A on K119
NMR: nuclear magnetic resonance
SSP: secondary structure propensity

## SUPPLEMENTARY FIGURES

**Supplemental Figure 1:**
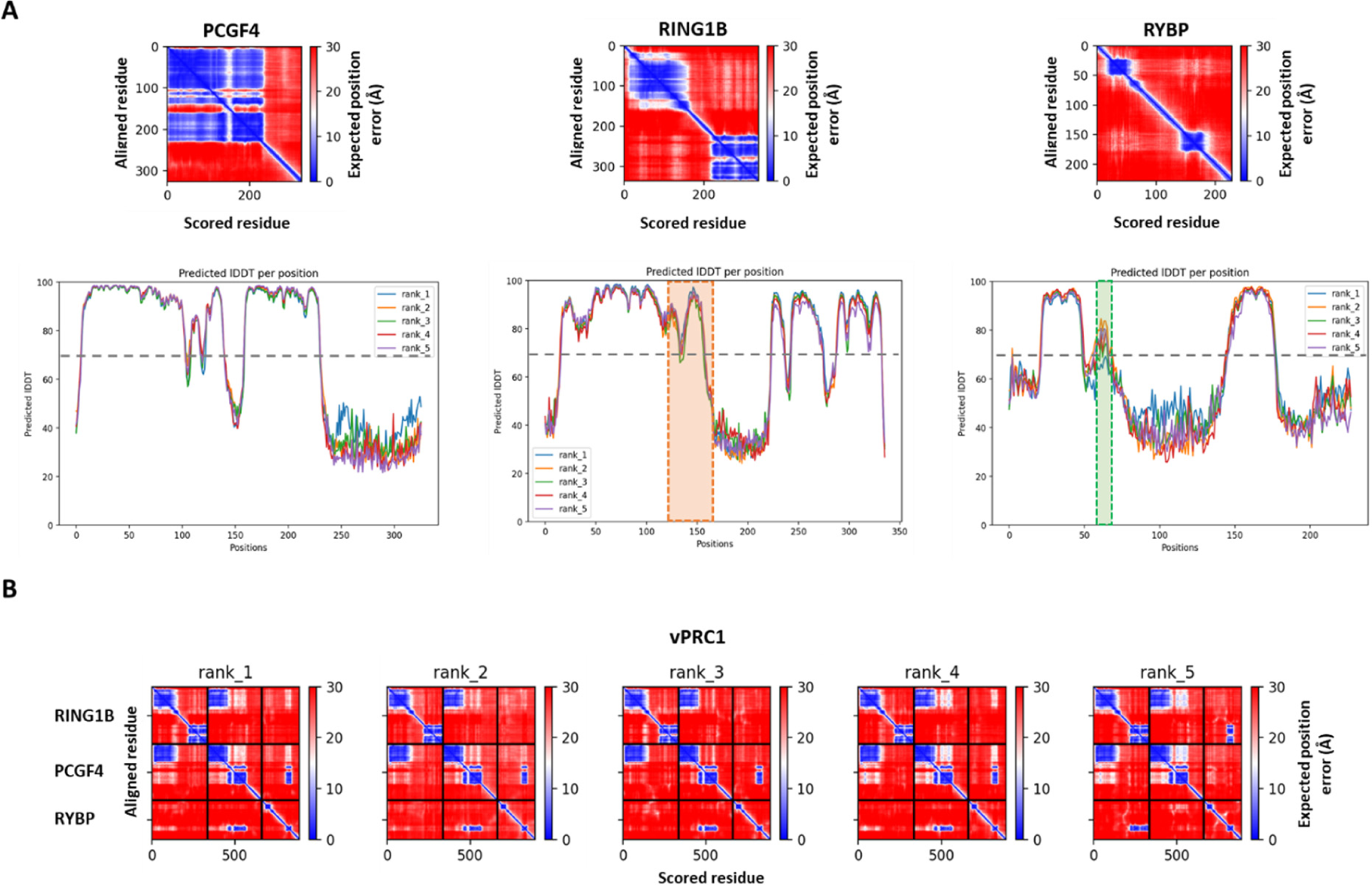
Statistics from AlphaFold2 models obtained for vPRC1.4 core complex. **A)** PAE and plDDT plots obtained for PCGF4, RING1B and RYBP subunits in isolation. The position of the newly identified structural motifs, RING1B ‘linker helix’ and RYBP ‘N-term helix’ is indicated in the plDDT plots by a dashed box, in orange and green, respectively. **B)** PAE plots obtained for the prediction models of the trimeric vPRC1.4 core complex.

**Supplemental Figure 2:**
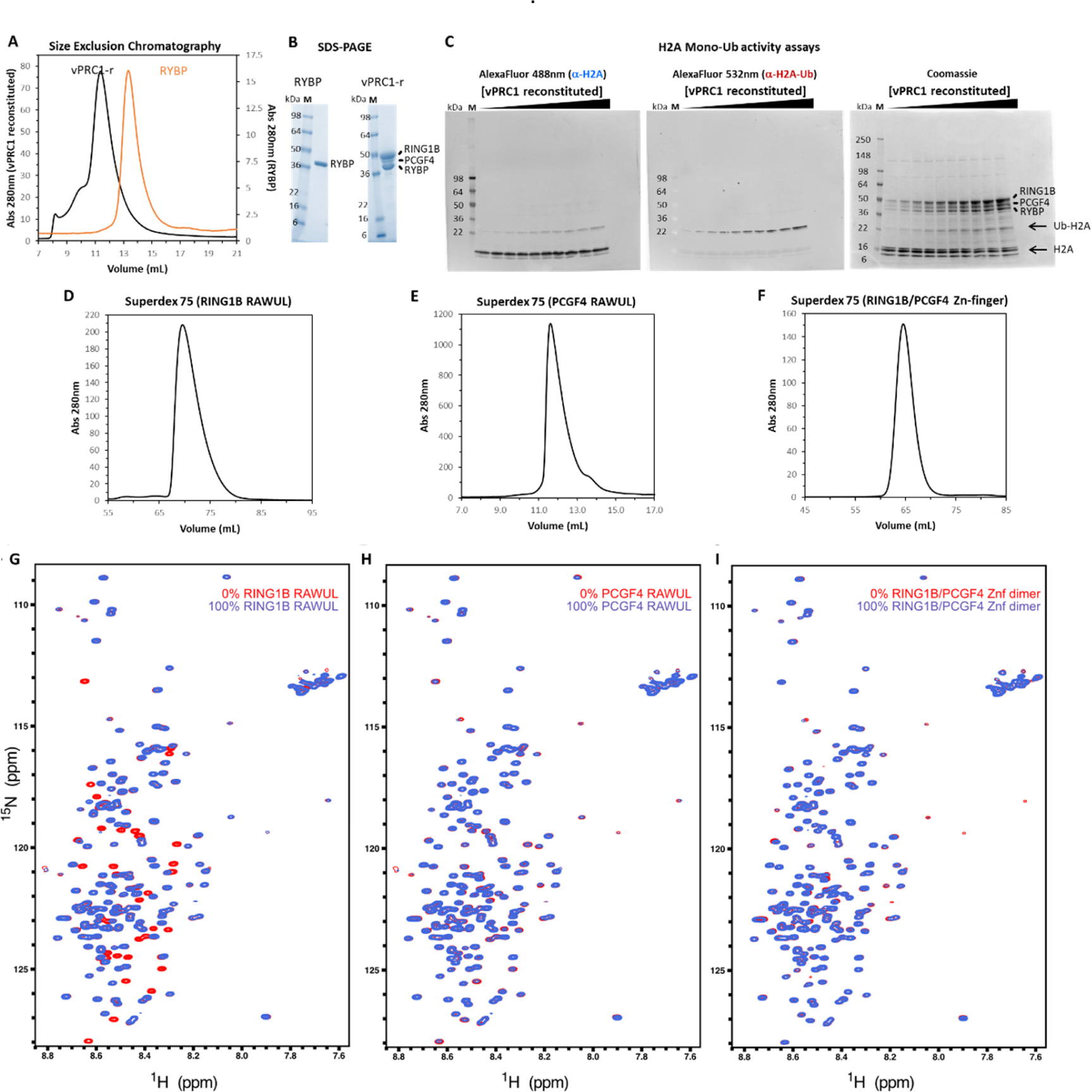
NMR analysis of RYBP binding to RING1B-PCGF4 catalytic heterodimer. **(A)** Size exclusion chromatography (SEC) elution profiles obtained for ^13^C/^15^N-labelled RYBP (orange) and reconstituted vPRC1.4 trimer (vPRC1-r; black) samples used for NMR experiments. **(B)** SDS-PAGE on the final samples obtained for ^13^C/^15^N-labelled RYBP and reconstituted vPRC1.4 trimer (vPRC1-r), after SEC. **(C)** The reconstituted vPRC1.4 complex is active, as shown by H2A monoubiquitination activity assays, revealed by western blot. **(D-F)** Size exclusion chromatography (SEC) elution profiles obtained for RING1B RAWUL **(D)**, PCGF4 RAWUL **(E)** and RING1B-PCGF4 Zn-finger dimer **(F)** domain samples isolated for the NMR titration experiments with ^15^N-labelled RYBP. **(G)** Full ^1^H-^15^N HSQC spectra of free RYBP (red) and with 100% (molar ratio) of RING1B RAWUL (blue). **(H)** Full ^1^H-^15^N HSQC spectra of free RYBP (red) and with 100% (molar ratio) of PCGF4 RAWUL (blue). **(I)** Full ^1^H-^15^N HSQC spectra of free RYBP (red) and with 100% (molar ratio) of RING1B/PCGF4 Zinc-finger dimer (blue).

**Supplemental Figure 3:**
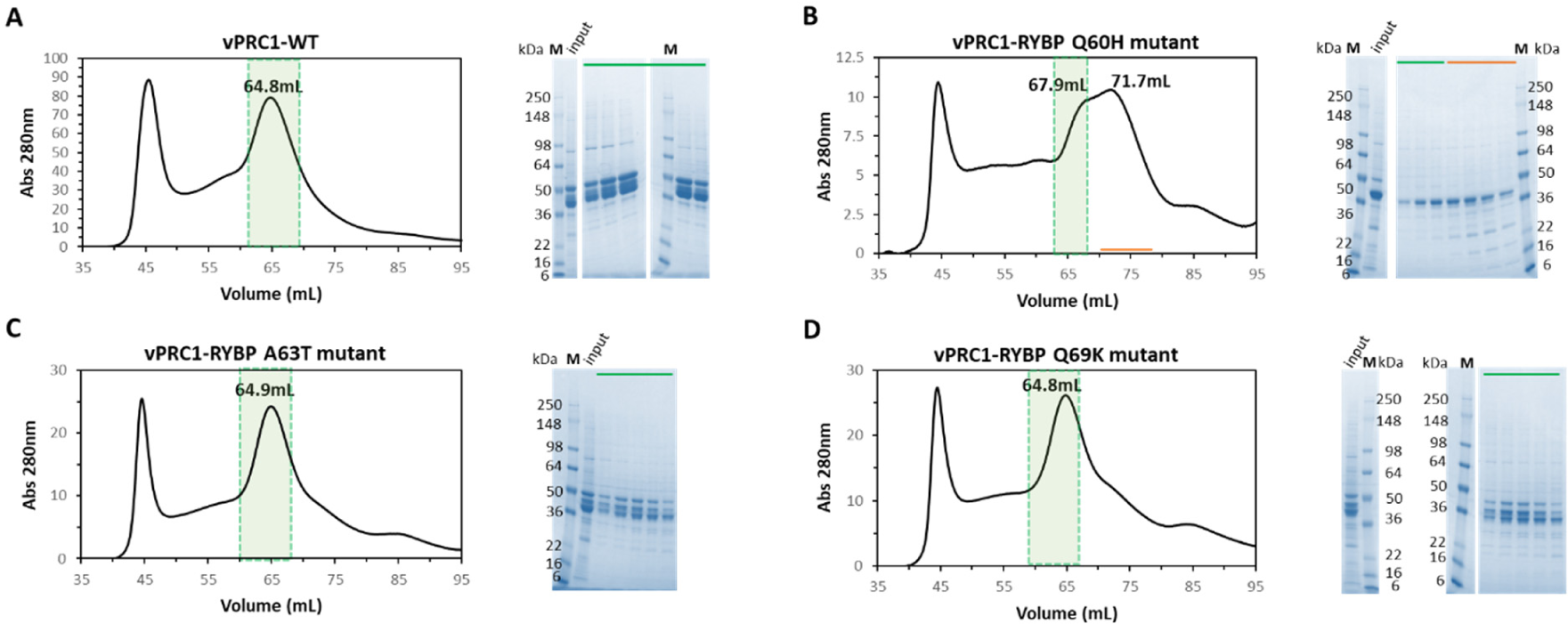
Purification of Q60H, A63T, and Q69K mutants. Size exclusion chromatography (SEC) elution profiles obtained for vPRC1 wt **(A)** and vPRC1-RYBP Q60H **(B)**, A63T **(C)** and Q69K **(D)** mutants, after Ni-Sepharose column, and corresponding SDS-PAGE gels on the indicated fractions. In green are highlighted the pooled fractions for pursuing of biophysical, biochemical and enzymatic characterizations as per Figure 7.

**Supplemental Figure 4:**
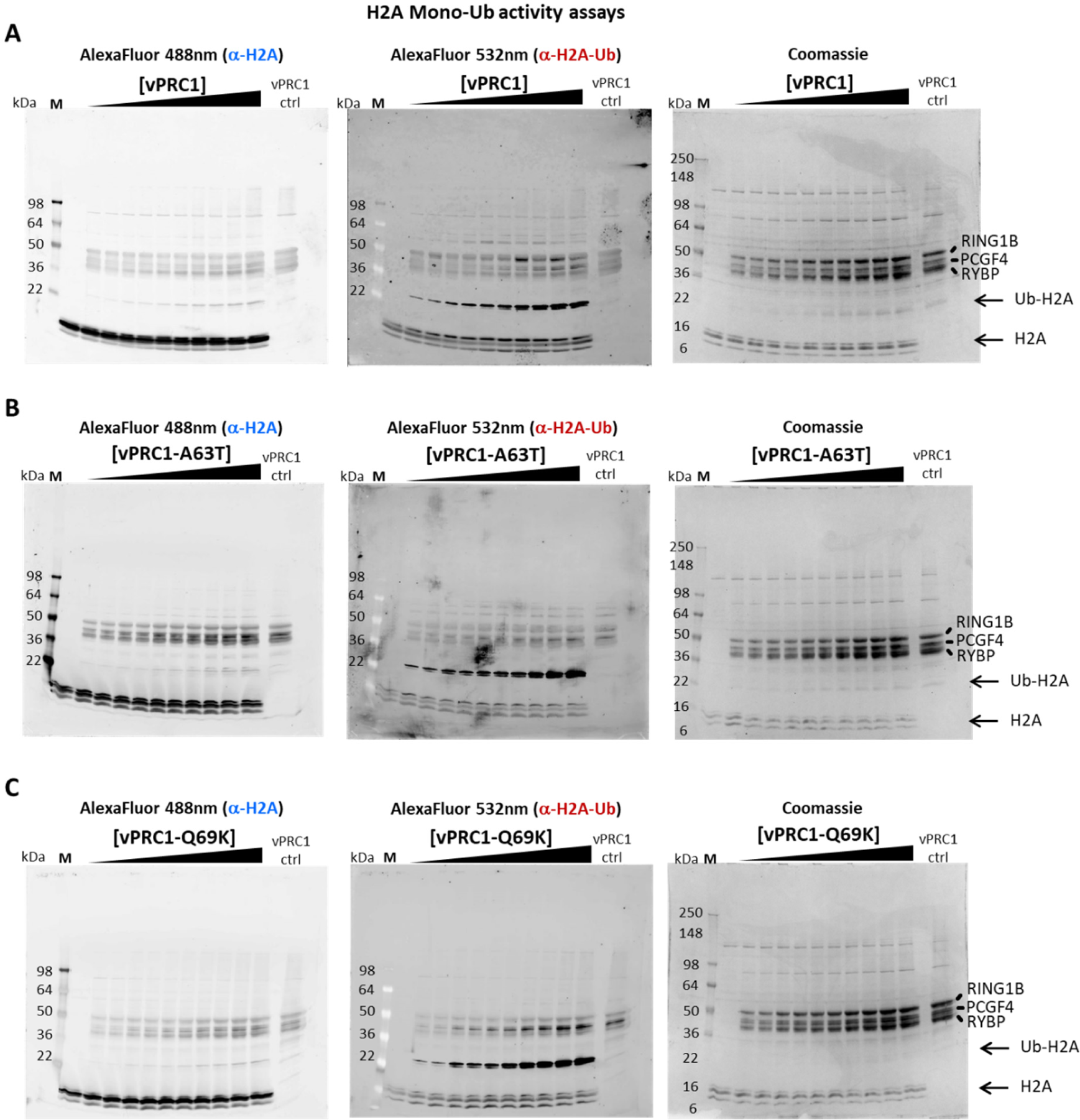
Enzymatic characterization of wt, A63T, and Q69K mutants. Examples of H2A monoubiquitination activity assays performed for vPRC1 wt **(A)** and vPRC1-RYBP A63T **(B)** and Q69K **(C)** mutants, as revealed by western blot. Experiments were repeated independently three times for each.

## SUPPLEMENTARY TABLES

**Supplementary Table 1:**
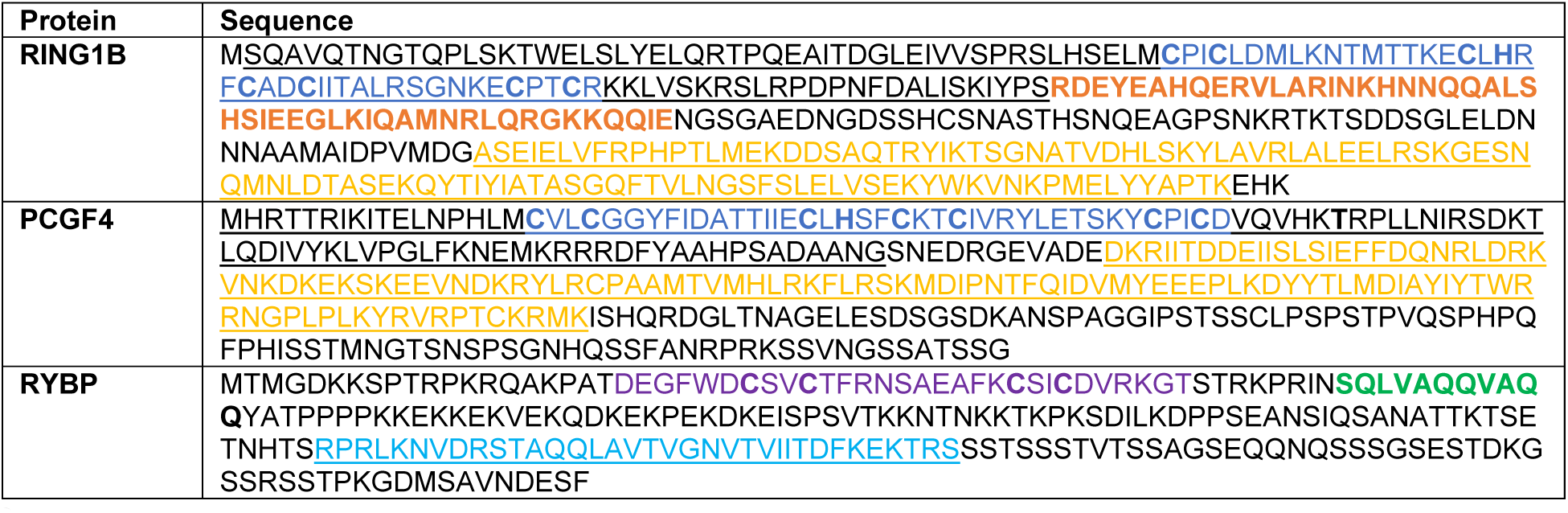
Protein sequences of human RING1B, PCGF4 and RYBP proteins used in this study for AlphaFold2 predictions. Sequence portions for which crystallographic structures have already been determined are underlined. For RING1B and PCGF4, RING and RAWUL domains are highlighted in dark blue and yellow, respectively. For RYBP, the Npl4 Zn-finger and β-hairpin are highlighted in purple and light blue respectively. The newly identified structural motifs on RING1B and RYBP are highlighted in bold, in orange and green, respectively.

## Notes

### Competing Interest Statement

The authors have declared no competing interest.

## REFERENCES

Alderson, T.R., Pritišanac, I., Kolarić, Đ., Moses, A.M., and Forman-Kay, J.D. (2023). Systematic identification of conditionally folded intrinsically disordered regions by AlphaFold2. bioRxiv.

Aloia, L., Di Stefano, B., and Di Croce, L. (2013). Polycomb complexes in stem cells and embryonic development. Development 140, 2525–2534.

Alshehri, M.A., Manee, M.M., Al-Fageeh, M.B., and Al-Shomrani, B.M. (2020). Genomic Analysis of Intrinsically Disordered Proteins in the Genus Camelus. Int J Mol Sci 21.

Arrigoni, R., Alam, S.L., Wamstad, J.A., Bardwell, V.J., Sundquist, W.I., and Schreiber-Agus, N. (2006). The Polycomb-associated protein Rybp is a ubiquitin binding protein. FEBS Lett 580, 6233–6241.

Bajusz, I., Kovács, G., and Pirity, M. (2018). From Flies to Mice: The Emerging Role of Non-Canonical PRC1 Members in Mammalian Development. Epigenomes 2.

Bejarano, F., Gonzalez, I., Vidal, M., and Busturia, A. (2005). The Drosophila RYBP gene functions as a Polycomb-dependent transcriptional repressor. Mech Dev 122, 1118–1129.

Bieniossek, C., Imasaki, T., Takagi, Y., and Berger, I. (2012). MultiBac: expanding the research toolbox for multiprotein complexes. Trends Biochem Sci 37, 49–57.

Blackledge, N.P., Farcas, A.M., Kondo, T., King, H.W., McGouran, J.F., Hanssen, L.L.P., Ito, S., Cooper, S., Kondo, K., Koseki, Y., et al. (2014). Variant PRC1 complex-dependent H2A ubiquitylation drives PRC2 recruitment and polycomb domain formation. Cell 157, 1445–1459.

Boeri Erba, E., Signor, L., Oliva, M.F., Hans, F., and Petosa, C. (2018). Characterizing Intact Macromolecular Complexes Using Native Mass Spectrometry. Methods Mol Biol 1764, 133–151.

Boeri Erba, E., Signor, L., and Petosa, C. (2020). Exploring the structure and dynamics of macromolecular complexes by native mass spectrometry. J Proteomics 222, 103799.

Boivin, S., Kozak, S., and Meijers, R. (2013). Optimization of protein purification and characterization using Thermofluor screens. Protein Expression and Purification 91, 192–206.

Camacho-Zarco, A.R., Schnapka, V., Guseva, S., Abyzov, A., Adamski, W., Milles, S., Jensen, M.R., Zidek, L., Salvi, N., and Blackledge, M. (2022). NMR Provides Unique Insight into the Functional Dynamics and Interactions of Intrinsically Disordered Proteins. Chemical Reviews 122, 9331–9356.

Chan, H.L., and Morey, L. (2019). Emerging Roles for Polycomb-Group Proteins in Stem Cells and Cancer. Trends Biochem Sci 44, 688–700.

Colombo, M., Pessey, O., and Marcia, M. (2019). Topology and enzymatic properties of a canonical Polycomb repressive complex 1 isoform. FEBS Lett 593, 1837–1848.

Cooper, S., Dienstbier, M., Hassan, R., Schermelleh, L., Sharif, J., Blackledge, N.P., De Marco, V., Elderkin, S., Koseki, H., Klose, R., et al. (2014). Targeting polycomb to pericentric heterochromatin in embryonic stem cells reveals a role for H2AK119u1 in PRC2 recruitment. Cell Rep 7, 1456–1470.

Delaforge, E., Kragelj, J., Tengo, L., Palencia, A., Milles, S., Bouvignies, G., Salvi, N., Blackledge, M., and Jensen, M.R. (2018). Deciphering the Dynamic Interaction Profile of an Intrinsically Disordered Protein by NMR Exchange Spectroscopy. J Am Chem Soc 140, 1148–1158.

Endoh, M., Endo, T.A., Shinga, J., Hayashi, K., Farcas, A., Ma, K.W., Ito, S., Sharif, J., Endoh, T., Onaga, N., et al. (2017). PCGF6-PRC1 suppresses premature differentiation of mouse embryonic stem cells by regulating germ cell-related genes. Elife 6.

Flavahan, W.A., Gaskell, E., and Bernstein, B.E. (2017). Epigenetic plasticity and the hallmarks of cancer. Science 357.

Fursova, N.A., Blackledge, N.P., Nakayama, M., Ito, S., Koseki, Y., Farcas, A.M., King, H.W., Koseki, H., and Klose, R.J. (2019). Synergy between Variant PRC1 Complexes Defines Polycomb-Mediated Gene Repression. Mol Cell 74, 1020–1036 e1028.

Gao, Z., Zhang, J., Bonasio, R., Strino, F., Sawai, A., Parisi, F., Kluger, Y., and Reinberg, D. (2012). PCGF homologs, CBX proteins, and RYBP define functionally distinct PRC1 family complexes. Mol Cell 45, 344–356.

Gray, F., Cho, H.J., Shukla, S., He, S., Harris, A., Boytsov, B., Jaremko, L., Jaremko, M., Demeler, B., Lawlor, E.R., et al. (2016). BMI1 regulates PRC1 architecture and activity through homo- and hetero-oligomerization. Nat Commun 7, 13343.

Haffke, M., Viola, C., Nie, Y., and Berger, I. (2013). Tandem recombineering by SLIC cloning and Cre-LoxP fusion to generate multigene expression constructs for protein complex research. Methods Mol Biol 1073, 131–140.

Hart, D.J., and Tarendeau, F. (2006). Combinatorial library approaches for improving soluble protein expression in Escherichia coli. Acta Crystallogr D Biol Crystallogr 62, 19–26.

Healy, E., and Bracken, A.P. (2020). If You Like It Then You Shoulda Put Two “RINGs” on It: Delineating the Roles of vPRC1 and cPRC1. Mol Cell 77, 685–687.

Jensen, M.R., Communie, G., Ribeiro, E.A., Jr., Martinez, N., Desfosses, A., Salmon, L., Mollica, L., Gabel, F., Jamin, M., Longhi, S., et al. (2011). Intrinsic disorder in measles virus nucleocapsids. Proc Natl Acad Sci U S A 108, 9839–9844.

Jumper, J., Evans, R., Pritzel, A., Green, T., Figurnov, M., Ronneberger, O., Tunyasuvunakool, K., Bates, R., Zidek, A., Potapenko, A., et al. (2021). Highly accurate protein structure prediction with AlphaFold. Nature 596, 583–589.

Junco, S.E., Wang, R., Gaipa, J.C., Taylor, A.B., Schirf, V., Gearhart, M.D., Bardwell, V.J., Demeler, B., Hart, P.J., and Kim, C.A. (2013). Structure of the polycomb group protein PCGF1 in complex with BCOR reveals basis for binding selectivity of PCGF homologs. Structure 21, 665–671.

Ko, K.T., Lennartz, F., Mekhaiel, D., Guloglu, B., Marini, A., Deuker, D.J., Long, C.A., Jore, M.M., Miura, K., Biswas, S., et al. (2022). Structure of the malaria vaccine candidate Pfs48/45 and its recognition by transmission blocking antibodies. Nat Commun 13, 5603.

Konarev, P.V., Volkov, V.V., Sokolova, A.V., Koch, M.H.J., and Svergun, D.I. (2003). PRIMUS: a Windows PC-based system for small-angle scattering data analysis. Journal of Applied Crystallography 36, 1277–1282.

Lakomek, N.A., Ying, J., and Bax, A. (2012). Measurement of (1)(5)N relaxation rates in perdeuterated proteins by TROSY-based methods. J Biomol NMR 53, 209–221.

Lau, M.S., Schwartz, M.G., Kundu, S., Savol, A.J., Wang, P.I., Marr, S.K., Grau, D.J., Schorderet, P., Sadreyev, R.I., Tabin, C.J., et al. (2017). Mutation of a nucleosome compaction region disrupts Polycomb-mediated axial patterning. Science 355, 1081–1084.

Leitner, A., Walzthoeni, T., and Aebersold, R. (2014). Lysine-specific chemical cross-linking of protein complexes and identification of cross-linking sites using LC-MS/MS and the xQuest/xProphet software pipeline. Nat Protoc 9, 120–137.

Lowary, P.T., and Widom, J. (1998). New DNA sequence rules for high affinity binding to histone octamer and sequence-directed nucleosome positioning. J Mol Biol 276, 19–42.

Madeira, F., Pearce, M., Tivey, A.R.N., Basutkar, P., Lee, J., Edbali, O., Madhusoodanan, N., Kolesnikov, A., and Lopez, R. (2022). Search and sequence analysis tools services from EMBL-EBI in 2022. Nucleic Acids Res 50, W276–W279.

Marsh, J.A., Singh, V.K., Jia, Z., and Forman-Kay, J.D. (2006). Sensitivity of secondary structure propensities to sequence differences between alpha- and gamma-synuclein: implications for fibrillation. Protein Sci 15, 2795–2804.

McGinty, R.K., Makde, R.D., and Tan, S. (2016). Preparation, Crystallization, and Structure Determination of Chromatin Enzyme/Nucleosome Complexes. Methods Enzymol 573, 43–65.

Mirdita, M., Schutze, K., Moriwaki, Y., Heo, L., Ovchinnikov, S., and Steinegger, M. (2022). ColabFold: making protein folding accessible to all. Nat Methods 19, 679–682.

Morgner, N., and Robinson, C.V. (2012). Massign: an assignment strategy for maximizing information from the mass spectra of heterogeneous protein assemblies. Anal Chem 84, 2939–2948.

Mosalaganti, S., Obarska-Kosinska, A., Siggel, M., Taniguchi, R., Turonova, B., Zimmerli, C.E., Buczak, K., Schmidt, F.H., Margiotta, E., Mackmull, M.T., et al. (2022). AI-based structure prediction empowers integrative structural analysis of human nuclear pores. Science 376, eabm9506.

Neira, J.L., Roman-Trufero, M., Contreras, L.M., Prieto, J., Singh, G., Barrera, F.N., Renart, M.L., and Vidal, M. (2009). The transcriptional repressor RYBP is a natively unfolded protein which folds upon binding to DNA. Biochemistry 48, 1348–1360.

Pasini, D., and Di Croce, L. (2016). Emerging roles for Polycomb proteins in cancer. Curr Opin Genet Dev 36, 50–58.

Pierce, S.B., Stewart, M.D., Gulsuner, S., Walsh, T., Dhall, A., McClellan, J.M., Klevit, R.E., and King, M.-C. (2018). De novo mutation in RING1 with epigenetic effects on neurodevelopment. Proceedings of the National Academy of Sciences 115, 1558–1563.

Robert, X., and Gouet, P. (2014). Deciphering key features in protein structures with the new ENDscript server. Nucleic Acids Res 42, W320–324.

Rose, N.R., King, H.W., Blackledge, N.P., Fursova, N.A., Ember, K.J., Fischer, R., Kessler, B.M., and Klose, R.J. (2016). RYBP stimulates PRC1 to shape chromatin-based communication between Polycomb repressive complexes. Elife 5.

Schneider, R., Blackledge, M., and Jensen, M.R. (2019). Elucidating binding mechanisms and dynamics of intrinsically disordered protein complexes using NMR spectroscopy. Curr Opin Struct Biol 54, 10–18.

Schuck, P. (2000). Size-distribution analysis of macromolecules by sedimentation velocity ultracentrifugation and lamm equation modeling. Biophys J 78, 1606–1619.

Schuettengruber, B., Bourbon, H.M., Di Croce, L., and Cavalli, G. (2017). Genome Regulation by Polycomb and Trithorax: 70 Years and Counting. Cell 171, 34–57.

Shim, Y., Duan, M.R., Chen, X., Smerdon, M.J., and Min, J.H. (2012). Polycistronic coexpression and nondenaturing purification of histone octamers. Anal Biochem 427, 190–192.

Simoes da Silva, C.J., Simon, R., and Busturia, A. (2018). Epigenetic and non-epigenetic functions of the RYBP protein in development and disease. Mech Ageing Dev 174, 111–120.

Simon, J.A., and Kingston, R.E. (2009). Mechanisms of polycomb gene silencing: knowns and unknowns. Nat Rev Mol Cell Biol 10, 697–708.

Sobott, F., Hernandez, H., McCammon, M.G., Tito, M.A., and Robinson, C.V. (2002). A tandem mass spectrometer for improved transmission and analysis of large macromolecular assemblies. Anal Chem 74, 1402–1407.

Solyom, Z., Schwarten, M., Geist, L., Konrat, R., Willbold, D., and Brutscher, B. (2013). BEST-TROSY experiments for time-efficient sequential resonance assignment of large disordered proteins. J Biomol NMR 55, 311–321.

Taherbhoy, A.M., Huang, O.W., and Cochran, A.G. (2015). BMI1-RING1B is an autoinhibited RING E3 ubiquitin ligase. Nat Commun 6, 7621.

Tavares, L., Dimitrova, E., Oxley, D., Webster, J., Poot, R., Demmers, J., Bezstarosti, K., Taylor, S., Ura, H., Koide, H., et al. (2012). RYBP-PRC1 complexes mediate H2A ubiquitylation at *Polycomb* target sites independently of PRC2 and H3K27me3. Cell 148, 664–678.

Team, R.C. (2013). R: A Language and Environment for Statistical Computing.

Trowitzsch, S., Bieniossek, C., Nie, Y., Garzoni, F., and Berger, I. (2010). New baculovirus expression tools for recombinant protein complex production. J Struct Biol 172, 45–54.

van den Heuvel, R.H., van Duijn, E., Mazon, H., Synowsky, S.A., Lorenzen, K., Versluis, C., Brouns, S.J., Langridge, D., van der Oost, J., Hoyes, J., et al. (2006). Improving the performance of a quadrupole time-of-flight instrument for macromolecular mass spectrometry. Anal Chem 78, 7473–7483.

Vijayachandran, L.S., Viola, C., Garzoni, F., Trowitzsch, S., Bieniossek, C., Chaillet, M., Schaffitzel, C., Busso, D., Romier, C., Poterszman, A., et al. (2011). Robots, pipelines, polyproteins: enabling multiprotein expression in prokaryotic and eukaryotic cells. J Struct Biol 175, 198–208.

Wang, L., Brown, J.L., Cao, R., Zhang, Y., Kassis, J.A., and Jones, R.S. (2004). Hierarchical recruitment of polycomb group silencing complexes. Mol Cell 14, 637–646.

Wang, R., Taylor, A.B., Leal, B.Z., Chadwell, L.V., Ilangovan, U., Robinson, A.K., Schirf, V., Hart, P.J., Lafer, E.M., Demeler, B., et al. (2010). *Polycomb* group targeting through different binding partners of RING1B C-terminal domain. Structure 18, 966–975.

Xia, Y., Ritz, C., Baty, F., Streibig, J.C., and Gerhard, D. (2015). Dose-Response Analysis Using R. Plos One 10.

